# FGFR3 is a positive regulator of osteoblast expansion and differentiation during zebrafish skull vault development

**DOI:** 10.1101/2020.01.02.884155

**Authors:** Emilie Dambroise, Ivan Ktorza, Alessandro Brombin, Ghaith Abdessalem, Joanne Edouard, Marine Luka, Imke Fiedler, Olivia Binder, Olivier Pelle, E. Elizabeth Patton, Björn Busse, Mickaël Menager, Frederic Sohm, Laurence Legeai-Mallet

## Abstract

Gain- or loss-of-function mutations in fibroblast growth factor receptor 3 (FGFR3) result in cranial vault defects - highlighting the protein’s role in membranous ossification. Zebrafish express high levels of *fgfr3* during skull development; in order to study FGFR3’s role in cranial vault development, we generated the first *fgfr3* loss-of-function zebrafish *(fgfr3^lof/lof^*). The mutant fish exhibited major changes in the craniofacial skeleton, with a lack of sutures, abnormal frontal and parietal bones, and the presence of ectopic bones. Integrated analyses (*in vivo* imaging, and single-cell RNA sequencing of the osteoblast lineage) of zebrafish *fgfr3^lof/lof^* revealed a delay in osteoblast expansion and differentiation, together with changes in the extracellular matrix. These findings demonstrate that *fgfr3* is a positive regulator of osteogenesis. We hypothesize that changes in the extracellular matrix within growing bone impair cell-cell communication, mineralization, and new osteoblast recruitment.

## Introduction

Cranial vault anomalies occur in many fibroblast growth factor receptor (FGFR)-related disorders. The progressive development of the cranial vault is initiated during embryogenesis, and is completed during adulthood by intramembranous ossification (except for the supra-occipital bones). Cranial vault formation starts with the condensation of mesenchymal cells, which then differentiate into the osteoprogenitors and osteoblasts required for bone matrix synthesis. The frontal and parietal bones develop from ossification centers formed by the deposition of non-mineralized bone matrix. At the edge of developing bones, new osteoblasts continually differentiate and thus expand the ossification centers. When two bones meet, a suture composed of mesenchymal tissue forms between two membranous zones.

The genes coding for members of the FGFR family are known to be involved in cranial vault formation. Along with the fibroblast growth factor (FGF) ligands, the FGFRs are expressed extensively in both developing and mature bones, where they regulate osteogenesis. Dysregulation of FGFRs is often associated with premature suture fusion; in the skull, this leads to craniofacial anomalies referred to as craniosynostoses (Wilkie et al., 2017). The p.Pro250Arg and p.Ala391Glu gain of function mutations in FGFR3 respectively cause Muenke syndrome (the most common craniosynostosis) and Crouzon syndrome with acanthosis nigricans – both of which are characterized by premature suture fusion in the skull vault (Meyers et al., 1995; Muenke et al., 1997). In earlier research, we observed a defect in membranous ossification (revealed by the presence of a large fontanelle) in achondroplasia (the most common form of dwarfism, due to a p.Gly380Arg FGFR3 gain-of-function mutation), (Di Rocco et al., 2014). Similarly, the absence of FGFR3 also affects skull formation, as demonstrated by the presence of microcephaly and Wormian bones in people with camptodactyly-tall stature-scoliosis-hearing loss syndrome (CATSHL) (Makrythanasis et al., 2014; Toydemir et al., 2006).

We previously described the partial, premature fusion of coronal sutures and impaired frontal bone formation in an (*Fgfr3^Y367C/+^*) murine model mimicking achondroplasia; this provided clear evidence that FGFR3 signaling is important for membranous ossification (Di Rocco et al., 2014). Surprisingly, the murine model of Muenke syndrome (*Fgfr3^P244R/+^*) rarely displays premature fusion of the coronal sutures, and the skull deformities are very mild (Twigg et al 2009). The *Fgfr3* knock-out mouse model does not have an obvious skull vault phenotype (Deng et al., 1996; Valverde-Franco et al., 2004). Hence, FGFR3’s specific role in skull vault formation has yet not been determined. In the zebrafish, Fgfr3 is highly conserved, and is expressed in mature osteoblasts within the osteogenic fronts and along the frontal bone (Topczewska et al 2016, Ledwon 2018). The late development and accessibility of the cranial vault in the juvenile zebrafish provides a unique opportunity to study each of the steps in osteogenesis during skull bone formation (Cornille et al., 2019; Topczewska et al., 2016). In the present report, we describe strong skull vault anomalies in an Fgfr3 loss-of-function mutant zebrafish model (*fgfr3^lof/lof^*) with abnormal frontal and parietal bones, the presence of ectopic bones, and a lack of suture formation. After we had used the transgenic fluorescent line *Tg*(*sp7*:*mCherry*; *bglap*:*GFP*) to label immature and mature osteoblasts, the sequential imaging of live zebrafish throughout their development showed that the differentiation and expansion of immature osteoblast were impaired *in fgfr3^lof/lof^* fish. Lastly, we used single-cell RNA sequencing to assess the impact of *fgfr3* loss of function on fish cranial vaults at 9 mm standard length (SL). We confirmed the presence of a defect in immature osteoblast differentiation, and observed marked differences in the expression of extracellular matrix (ECM) components. We hypothesized that in the absence of Fgfr3, defects in the ECM would affect cell-cell communication and the recruitment of new osteoblasts during bone growth. We conclude that Fgfr3 is a positive regulator of osteogenesis during cranial vault formation.

## Results

### Inhibition of Fgfr3’s tyrosine kinase activity affects cranial vault development in zebrafish

The zebrafish *fgfr3* gene is strongly transcribed in osteoblasts during cranial vault formation (Ledwon et al., 2018). The alignment of the deduced amino acid sequence of zebrafish Fgfr3 with that of its human homolog (FGFR3) shows a high level of homology between the two proteins (77% overall, and more than 90% when considering the tyrosine kinase domain alone; Figure 1 - supplement 1). In order to investigate Fgfr3’s role, we used the tyrosine kinase inhibitor BGJ398 (infigratinib, LC Laboratories) to block the factor’s tyrosine kinase activity during cranial vault development (Gudernova et al., 2016; Komla-Ebri et al., 2016). Zebrafish larvae with a standard length (SL) of 7 mm (7SL) and expressing mCherry in osteoblasts (*Tg*(*sp7*: *mCherry*) were injected with BGJ398 every other day for 15 days (Figure 1A). After treatment, BGJ398-treated larvae had the same SL as the larvae treated with vehicle (3% DMSO), confirming that BGJ398 did not interfere with overall growth (Figure 1B). We found that the mean ± SD surface area of the fontanelle (i.e. the osteoblast-free area of the cranial vault) was significantly larger in BGJ398-treated larvae (347791 ± 50768 µm²) than in DMSO-treated larvae (135171 ± 22043 µm²; p=0.014) (Figure 1C). Our results indicate that in the zebrafish, the inhibition of Fgfr3 activity strongly disrupts osteoblast invasion during cranial vault development.

**Figure 1:**
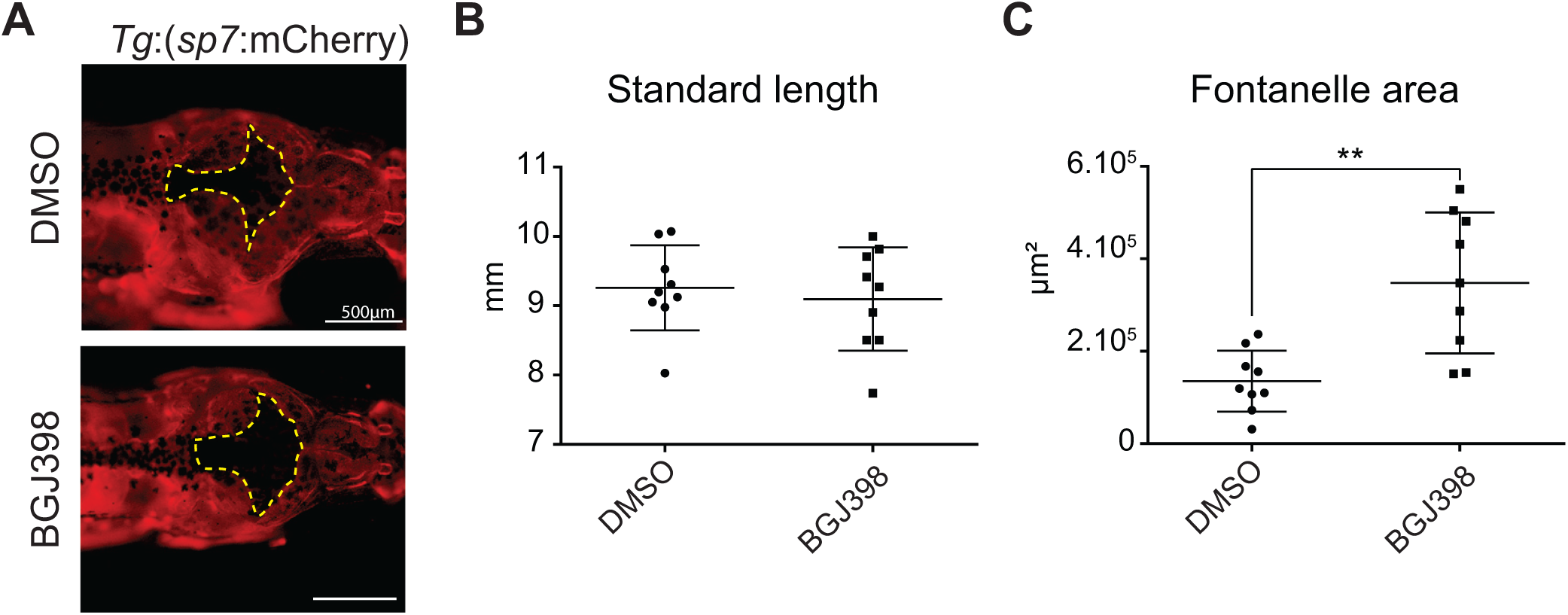
Inhibition of cranial vault development by BGJ398 injection. (A) Dorsal view of the *Tg*(*sp7*:*mCherry*) line injected with 3% DMSO or 2 mg/kg BGJ398. Dotted lines indicate the fontanelle area, with no mCherry-positive osteoblasts. (B) Standard length and (C) fontanelle area of fish treated with 3% DMSO (n=9) or BGJ398 (n=9) every other day for 15 days. P-values were determined using Student’s t-test; the error bars correspond to the standard error of the mean.

### *fgfr3* loss of function induces craniofacial and axial skeleton anomalies in three-month-old zebrafish

We next used CrispR/Cas9 technology to generate two different loss-of-function *fgfr3* zebrafish lines, with a stop codon at position 377 for *fgfr3^lof1/lof1^* (Figure 2A) and at position 278 for *fgfr3^lof2/lof2^* (Figure 2 supplement 1). The predicted sequence indicated that both mutations led to the formation of a truncated protein lacking both the transmembrane domain and the tyrosine kinase domain. Since the two mutations produced identical phenotypes, we conducted all subsequent experiments with *fgfr3^lof1/lof1^* line. A quantitative real-time PCR (qPCR) analysis confirmed that *fgfr3* expression was abnormally low in *fgfr3^lof1/lof1^* zebrafish, whereas the expression of *fgfr1a, fgfr1b* and *fgfr2* was not affected (Figure 2B).

**Figure 2:**
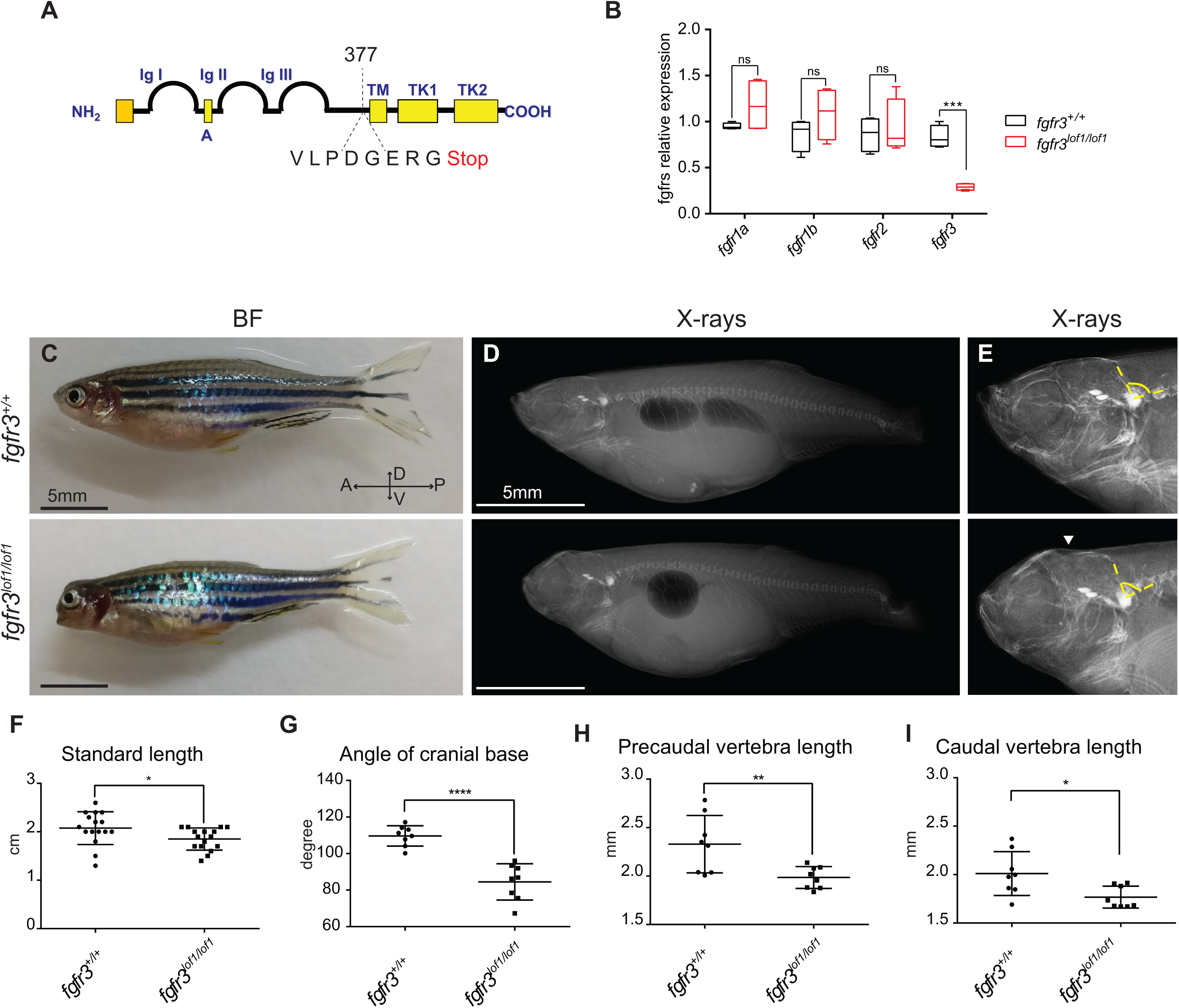
Three-month-old *fgfr3^lof1/lof1^* fish present craniofacial and axial skeletal anomalies. (A) Schematic representation of the Fgfr3 receptor, with the position of the lof1 mutation. IG: immunoglobulin domain; TM: transmembrane domain; TK: tyrosine kinase domain. (B) Relative expression of *fgfr* mRNA in 6-week-old *fgfr3^+/+^* (n=4) and *fgfr3^lof1/lof1^* fish (n=4). (C) Image of sibling *fgfr3^+/+^* and *fgfr3^lof1/lof1^* fish at 3 months of age; the *fgfr3^lof1/lof1^* fish present a flat face, microcephaly, and a low SL. (D) X-rays reveal severe anomalies of the craniofacial skeleton and the swim bladder in the *fgfr3^lof1/lof1^* mutant. (E) Magnified X-ray of the head in *fgfr3^+/+^* and *fgfr3^lof1/lof1^* fish, showing hypomineralization of the cranial vault (white arrow) and an abnormally small angle between the base of the cranium and the spine in *fgfr3^lof1/lof1^* fish (yellow dotted line). (F-I) Measurement of the SL (*fgfr3^+/+^* n=16; *fgfr3^lof1/lof1^* n=16), the cranial base angle (*fgfr3^+/+^* n=8; *fgfr3^lof1/lof1^* n=8), and the precaudal and caudal vertebra length (*fgfr3^+/+^* n=8; *fgfr3^lof1/lof1^* n=8). ns: non-significant; BF: bright field; A: anterior; P: posterior; D: dorsal; V: ventral. P-values were determined using Student’s t-test; the error bars correspond to the standard error of the mean.

Three-month-old *fgfr3^lof1/lof1^* fish displayed major craniofacial defects, with a flat face and microcephaly (Figure 2C). X-ray analyses confirmed that *fgfr3^lof1/lof1^* had an abnormal skull shape and that the cranial vault was less mineralized than in *fgfr3^+/+^* fish (Figure 2D, E). Moreover, *fgfr3^lof1/lof1^* fish were shorter than their control siblings (1.85 ± 0.06 cm vs. 2.07 ± 0.06 cm, respectively; p=0.035) (Figure 2F). *fgfr3^lof1/lof1^* fish also displayed a more strongly curved spine than controls, as measured by the angle formed between the base of the skull and the Weberian apparatus (*fgfr3*^+/+^:109.7 ± 2.0°; *fgfr3^lof1/lof1^*: 84.5 ± 3.5°; p<0.0001) (Figure 2D,G). *fgfr3^lof1/lof1^* fish also had shorter precaudal and caudal vertebrae than *fgfr3*^+/+^ fish, although the intervertebral spacing was unaffected (Figure 2 H, I and data Figure 2). Alcian blue and Alizarin red S staining demonstrated that *fgfr3^lof1/lof1^* fish had normally shaped fins (Figure 2 - supplement 2).

Lastly, we noticed that the swim bladder of *fgfr3^lof1/lof1^* was formed by a single chamber; this contrasted with the anterior chamber and posterior chamber that comprise the wild-type swim bladder (Figure 2C). These results indicate that the absence of Fgfr3 prevents evagination from the cranial end of the swim bladder’s original sac (Robertson et al., 2007).

The low SL, flat face, microcephaly, cranial vault hypomineralization and swim bladder defects were also observed in *fgfr3^lof2/lof2^* fish *-* confirming that these phenotypes are directly correlated with *fgfr3* loss-of-function mutations (Figure 2 supplement 2). Taken as a whole, our findings indicate that Fgfr3 has a role in development of the craniofacial and axial skeletons in zebrafish.

### *fgfr3* loss of function induces severe frontal and parietal bone defects, together with Wormian bone formation

To better characterize the craniofacial defects in *fgfr3^lof1/lof1^*, we used Alizarin red S staining and micro computed tomography (µCT) to study three-month-old fish. The µCT analyses confirmed that *fgfr3^lof1/lof1^* fish had a microcephalic skull, with low skull height and length (Figure 3A, B) and a normal skull width (Figure 3D-F). We observed that the foramen of the cranial vault was larger in *fgfr3* mutants than in control siblings. The mean ± SD surface area of the fontanelle (located between the frontal and parietal bones) was 0.0226 ± 0.0091 mm² in *fgfr3*^+/+^ fish and 0.988 ± 0.186 mm² in *fgfr3^lof1/lof1^* fish (p=0.0009; (Figure 3G-I). The abnormally large fontanelle in *fgfr3^lof1/lof1^* fish was associated with significantly smaller frontal bones (relative decrease: 42%) and parietal bones (relative decrease: 46%). The mean ± SD thickness of the parietal bone was not affected (*fgfr3*^+/+^:13.8 ± 0.7 µm; *fgfr3^lof1/lof1^*: 12.5 ± 1.4 µm; p=0.4251). Interestingly, we noted the presence of ectopic bones between the frontal and parietal bones in *fgfr3^lof1/lof1^* fish (Figure 3B, C). The ectopic bones resembled the Wormian bones described in patients carrying a homozygous *FGFR3* loss-of-function mutation (Makrythanasis et al., 2014). To study changes in the cranial vault phenotype during growth, we analyzed 6-month-old fish (Figure 3 supplement 1 A-C). Mutant fish had an irregularly shaped skull vault with a fontanelle still present, and 70% of the fish had ectopic bones (Figure 3 supplement 1 C).

**Figure 3:**
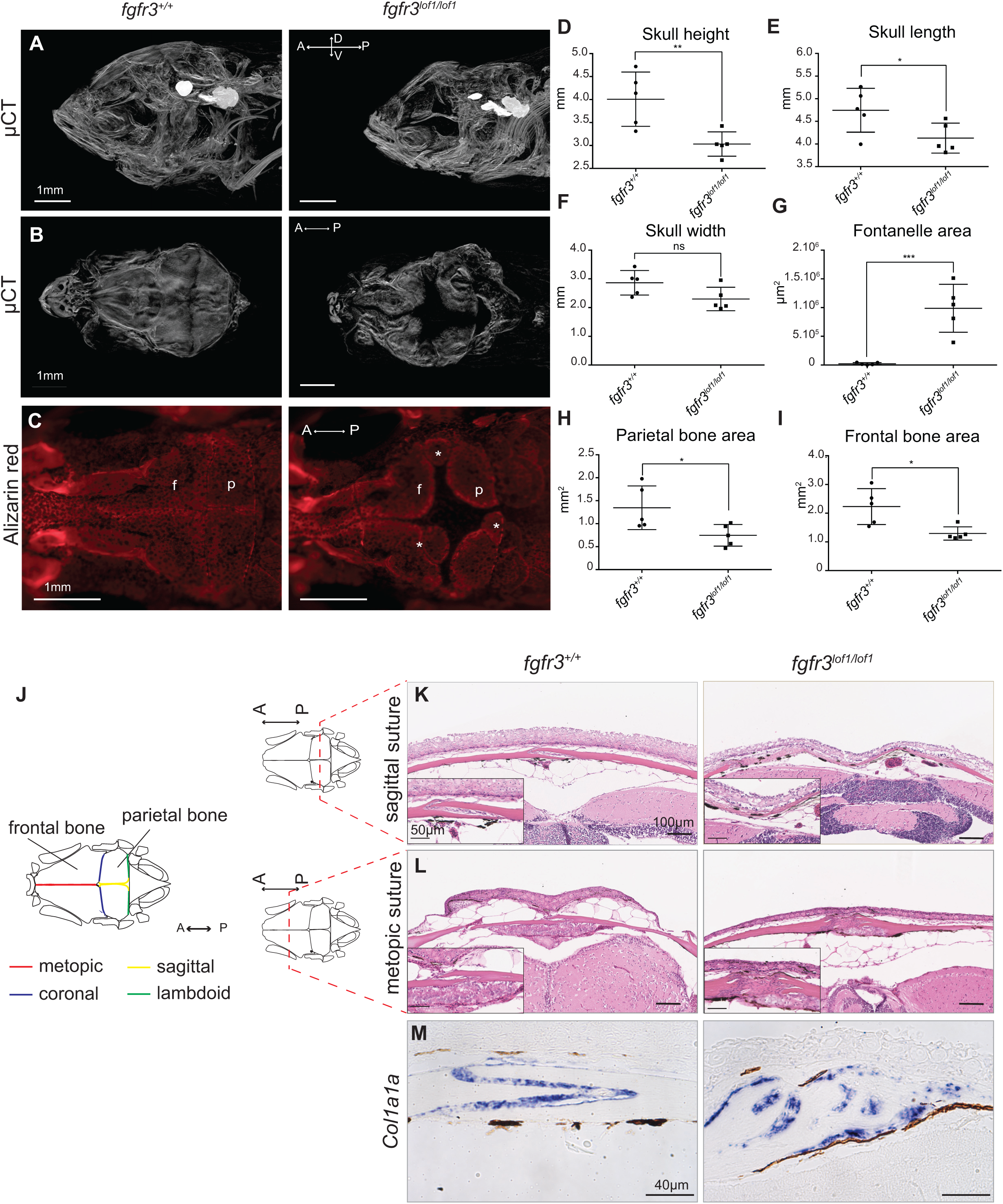
*fgfr3^lof1/lof1^* adult fish presented severe microcephaly, frontal and parietal bone defects, the presence of ectopic bones, and the absence of sutures. (A, B) The µCT profile and dorsal view of the craniofacial skeleton in 3-month-old *fgfr3^+/+^* and *fgfr3^lof1/lof1^* fish. (C) A dorsal view of the cranial vault stained with Alizarin red S highlighted frontal and parietal bone defects and the presence of ectopic bones (white star) in *fgfr3^+/+^* and *fgfr3^lof1/lof1^* mutants. (D-I) Measurement of skull height, length, and width, fontanelle area, and parietal and frontal bone areas (*fgfr3^+/+^* n=5; *fgfr3^lof1/lof1^* n=5). (J) Schematic representation of the zebrafish cranial vault. (K) Sections stained with hematoxylin-eosin reagent show the absence of sagittal sutures in *fgfr3^lof1/lof1^* fish. (L) Sections stained with hematoxylin-eosin reagent show the absence of metopic sutures in *fgfr3^lof1/lof1^* fish. (M) *In situ* hybridization reveals abnormal *col1a1a* expression (red arrows) at the osteogenic front in *fgfr3^lof1/lof1^* bones. ns: non-significant; A: anterior; P: posterior; D: dorsal; V: ventral; f: frontal; p: parietal. P-values were determined using Student’s t-test; the error bars correspond to the standard error of the mean.

We next investigated the morphology of the sutures in the cranial vault. Control fish displayed the typical sagittal and metopic sutures formed by the overlaps between the two parietal bones and the two frontal bones, respectively (Figure 3 K, L). The two osteogenic fronts overlapped but remained separated by a thin band of fibrous connective tissue that contained mesenchymal cells. Interestingly, we observed defective suture formation in *fgfr3^lof1/lof1^* fish. At the sagittal suture, the parietal bones’ osteogenic fronts were separated by a broad band of fibrous connective tissue (Figure 3 K). At the metopic suture, the frontal bones presented irregular osteogenic fronts separated by a large number of fibrotic cells (Figure 3 L). An analysis of *Col1a1a* expression (the gene coding for collagen type 1a1) confirmed the anomalies of the osteogenic front and revealed the presence of osteoblasts at osteogenic front in *fgfr3^lof1/lof1^* fish (Figure 3M). Nevertheless, cells expressing *Col1a1a* were visible in *fgfr3^lof1/lof1^* ectopic sites, such as the interdigital spaces formed by abnormal bones (red arrows). Metopic and sagittal sutures were still absent in the cranial vault of 6-month-old fish (Figure 3 supplement 1 D, E). The absence of suture formation in *fgfr3^lof1/lof1^* zebrafish shows that Fgfr3’s role in suture formation is conserved between zefrafish and humans.

With regard to the viscerocranium, we did not observe any obvious abnormalities in the Meckel cartilage of *fgfr3^lof1/lof1^* mutants (Figure 3, Table 1). Macroscopic analyses of *fgfr3*^lof1/lof1^ larvae on days 5 and 9 post-fertilization (dpf) did not reveal any developmental defects; there were no significant head abnormalities or delays in branchial arch ossification (Figure 3 supplement 2 A-C). The endochondral bones of the viscerocranium were not affected, as confirmed by measurements of the Meckel cartilage, palatoquadrate, and ceratohyal in *Tg(col2: mCherry) fgfr3^lof1/lof1^* and *fgfr3^+/+^* larvae 14 dpf (Figure 3 supplement 2 D-G).

Delayed parietal and frontal bone formation in *fgfr3^lof1/lof1^* fish was evidenced by the presence of large fontanelles. The ectopic bone formation described above appeared to compensate for this delay. The absence of viscerocranial anomalies suggests that the difference in skull shape was mainly due to bone abnormalities in the cranial vault.

### Osteoblast expansion and differentiation are affected *in fgfr3^lof1/lof1^* fish

We next used Alizarin red S staining to study the tissues that are mineralized during craniofacial skeleton formation from 8SL to 11SL (Figure 4A). From 9SL onwards, Alizarin red S staining of the neurocranium and the dermatocranium was slightly less intense in *fgfr3^lof1/lof1^* fish than in *fgfr3*^+/+^ fish. This phenotype developed progressively, as highlighted by our observations of 11SL fish. To assess the bone formation process in the cranial vault, we performed Alizarin red S *in vivo* labeling at 9SL and then calcein labeling 7 days later (Figure 4B). There were significantly less newly formed bones in *fgfr3^lof1/lof1^* fish (6 ±0.2 µm/day) than in *fgfr3*^+/+^ fish (26 ± 1.2 µm/day; p<0.0001) (Figure 4C). These results showed that Fgfr3 loss of function impaired the formation of cranial vault bones and had much the same effect as BGJ398 treatment (Figure 1A, 1C).

**Figure 4.**
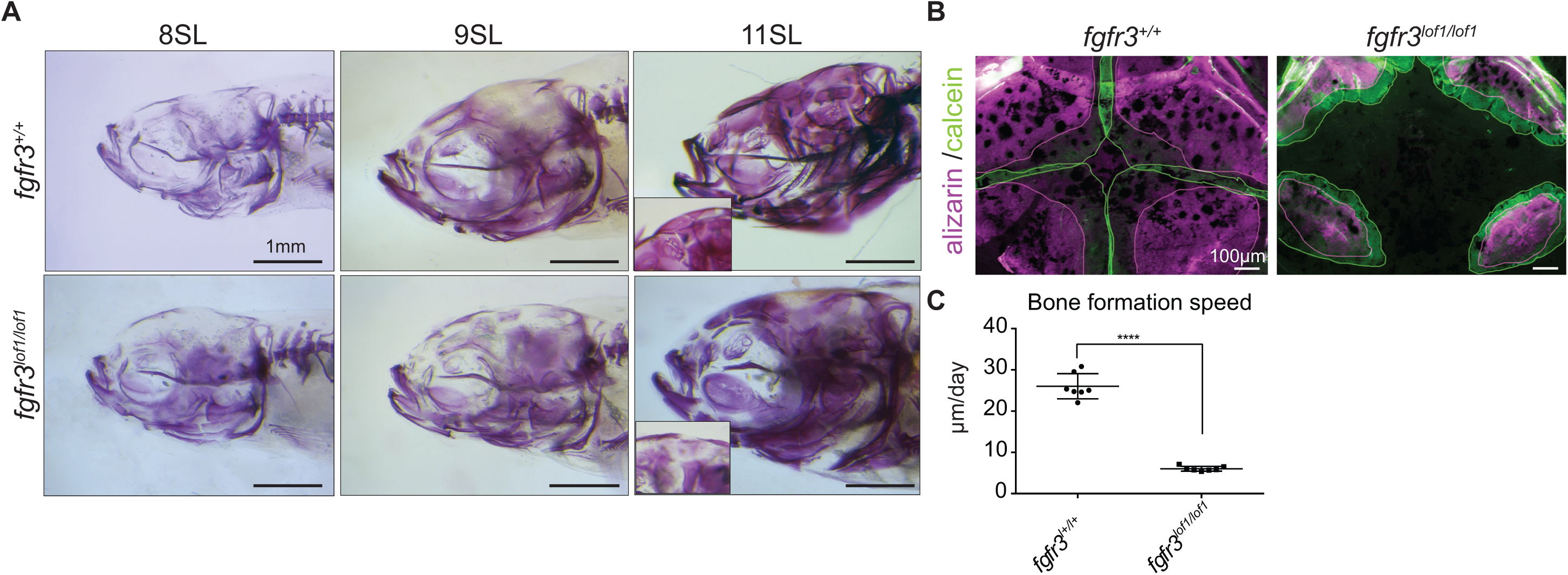
Cranial bone formation is delayed in *fgfr3^lof1/lof1^* fish. (A) Alizarin red S staining of *fgfr3^lof1/lof1^* and their siblings from 7 to 11mm SL; a high-magnification view shows the lack of cranial vault mineralization in *fgfr3^lof1/lof1^* fish at 11SL. (B) A dorsal view of the central skull vault region in *fgfr3^lof1/lof1^* fish and their siblings after Alizarin red S staining (in purple) at 9mm SL and then calcein staining (in green) 7 days later. (C) The bone formation rate in *fgfr3^lof1/lof1^* fish and their siblings. P-values were determined using Student’s t-test; the error bars correspond to the standard error of the mean.

Given that bones are formed by the action of osteoblasts, we decided to study these cells during cranial vault formation. We used *Tg*(*Sp7:mCherry; bglap:GFP*); *fgfr3*^+/+^/*fgfr3^lof1/lof1^* fish to express mCherry and GFP in immature and mature osteoblasts, respectively, with weekly sequential imaging of the fish from 7 (SL) to 11SL (Figure 5A). The ossification centers of the parietal and frontal bones were present at 7SL in *fgfr3^+/+^* fish (as revealed by the presence of immature osteoblasts) and expanded at later stages of development (Figure 5A). At 11SL, the immature osteoblasts of the frontal and parietal bones overlapped and formed coronal sutures. Mature osteoblasts were present in frontal bones at 7SL and in parietal bones at 8SL. At 9SL and 11SL, mature osteoblasts almost entirely covered the frontal and parietal bones; only a thin band of immature osteoblasts remained at the edges.

**Figure 5.**
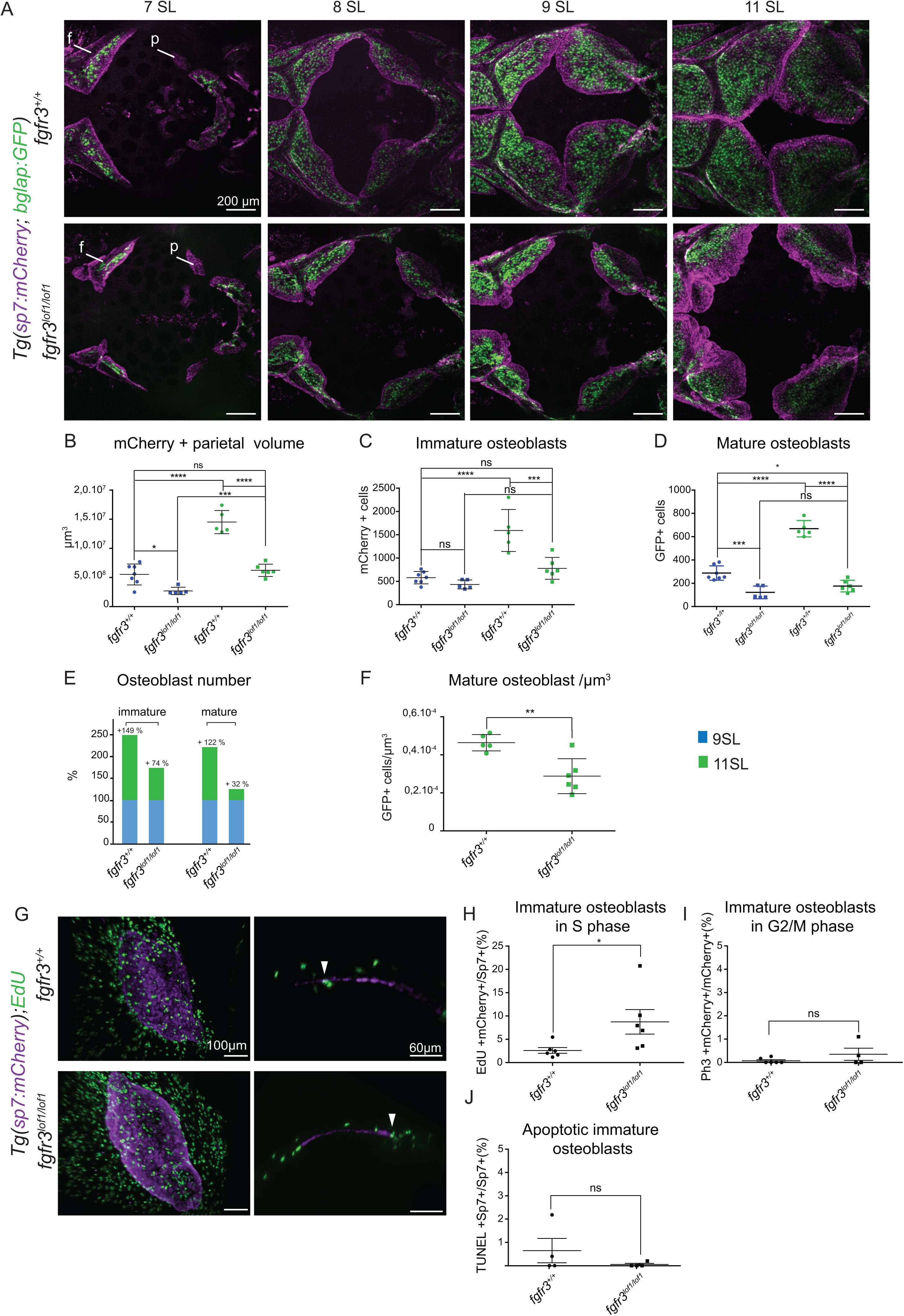
*fgfr3* loss of function impairs the proliferation and differentiation of immature osteoblasts. (A) Sequential weekly imaging of *Tg(sp7:mCherry; bglap:GFP)*; *fgfr3*^+/+^ and *Tg*(*sp7:mCherry;bglap:GFP*); *fgfr3^lof1/lof1^* fish from 7SL to 11SL. *fgfr3*^+/+^ immature osteoblasts (mCherry+) from parietal bone expanded from 7SL to 11SL to form the cranial vault, and then overlapped to form sutures. The differentiation of immature osteoblasts started at 7SL at the center of each bone. At 11SL, mature osteoblasts (GFP+) covered the entire cranial vault, with the exception of the bone formation front. *fgfr3^lof1/lof1^* immature osteoblasts failed to expand at 9SL, and a fall in the number of mature osteoblasts was observed. (B-D) The mCherry+ parietal volume and the immature (mCherry+) and mature (GFP+) osteoblast counts. (E) Immature and mature osteoblast counts from 9SL to 11SL (F) Quantification of mature osteoblasts/µm^3^ in parietal bones of 11SL *Tg*(*sp7:mCherry; bglap:GFP*); *fgfr3*^+/+^ and *Tg*(*sp7:mCherry; bglap:GFP*); *fgfr3*^lof1/lof1^ (, 11SL-*fgfr3^+/+^*: n= 5; 11 SL-*fgfr3^lof1/lof1^*: n=6) (G) Parietal bones stained with EdU and anti-mCherry in *Tg*(*sp7:mCherry*); *fgfr3* ^+/+^ and *Tg*(*sp7:mCherry*); *fgfr3^lof1/lof1^* fish. A cross-sectional view of the parietal bone shows mCherry- and EdU-positive cells (white arrows). (H) Quantification of the percentage of mCherry + cells that were EdU+ (*fgfr3^+/+^* n= 6; *fgfr3^lof1/lof1^* n=6). (I) The percentage of mCherry + and pH3+ cells (*fgfr3^+/+^* n= 5; *fgfr3^lof1/lof1^* n=4). (J) Quantification of TUNEL positive cells (*fgfr3^+/+^* n= 4; *fgfr3^lof1/lof1^* n=4). f: frontal; p: parietal; ns: non-significant, (B-D) A one-way ANOVA with Tukey’s multiple comparison test was performed to determine the statistical significance of inter-sample differences in the fold-change in each category of fish.(F-H) P-values were determined using Student’s t-test the error bars correspond to the standard error of the mean.

Interestingly, we found that immature osteoblasts were present in both *fgfr3^lof1/lof1^* and control fish at 7SL; however, the mutant’s frontal bones seemed to be fragmented (Figure 5A). From 8SL onwards, there were smaller ossification centers in *fgfr3^lof1/lof1^* fish than in controls (Figure 5A, 5B); this difference was accentuated between 9SL and 11SL and was associated with an irregular bone formation front. Quantitative analyses of *fgfr3^lof1/lof1^* and *fgfr3*^+/+^ parietal bones at 9SL and 11SL showed that as expected, a low parietal volume was correlated with a low immature osteoblast (mCherry^+^) count (Figure 5B, 5C). Comparisons of immature osteoblast counts on parietal bones from 9SL to 11SL revealed an impairment in cell expansion in *fgfr3^lof1/lof1^* fish; the number of immature osteoblasts increased by 149% in wild-type fish but only by 74% in *fgfr3^lof1/lof1^* fish (Figure 5E). Moreover, the defective expansion of immature osteoblasts was associated with a low mature osteoblast count at 9SL and 11SL (Figure 5A, 5D). The mature osteoblast count increased by 122% from 9SL to 11SL in wild-type fish but only by 32% in the mutant. This was consistent with the huge surface area covered with only immature osteoblasts at the bone formation front at 11SL (Figure 5A, 5E). The presence of a maturation defect was confirmed by the abnormally low number of GFP-positive cells per µm^3^ at 11SL in *fgfr3^lof1/lof1^* fish, relative to *fgfr3^+/+^* fish (Figure 5F). These findings demonstrated that the osteoblast maturation is dramatically impaired in *fgfr3^lof1/lof1^* zebrafish.

We next used EdU incorporation and anti-pH3 immunolabeling to study the proliferation of immature osteoblasts (Figure 5G) at 9SL. Most of the EdU^+^ mCherry^+^ cells in parietal bones were present at the edges of the ossification centers in both controls and mutants. However, we observed a significantly greater number of EdU^+^ mCherry^+^ cells in *fgfr3^lof1/lof1^* fish (Figure 5H). Following up on these findings, we observed that the mCherry^+^ cell count in G2/M (using an anti-pH3 antibody) was similar in mutants and controls (Figure 5I). Hence, the absence of Fgfr3 appeared to lead to dysregulation of the immature osteoblast cell cycle, with a longer S phase in mutants than in controls. This relative increase in the cell cycle duration was consistent with the delay in the immature osteoblast expansion observed between 9SL and 11SL in the mutant. Lastly, we investigated the apoptosis of the immature osteoblasts; TUNEL analyses revealed that the impaired bone formation was not due to premature death of the immature osteoblasts (Figure 5J).

In conclusion, our results demonstrate that Fgfr3 is a positive regulator of immature osteoblast expansion and differentiation during cranial vault formation. Nevertheless, the major delay in ossification center formation could not be explained by the observed changes of their cell cycle alone. Our findings indicated that the absence of Fgfr3 also affected the recruitment of newly immature osteoblasts formed from osteoprogenitors.

### Identification of osteogenic cell subpopulations involved in cranial vault formation with single-cell RNA sequencing

In order to better characterize the cellular impact of *fgfr3* loss of function, we performed a single-cell RNA sequencing on *fgfr3^lof1/lof1^* and *fgfr3^+/+^* cranial vaults at 9SL. We used the Chromium system (10x Genomics) to profile the transcriptome of nearly 28000 single cells isolated from cranial vaults from two *fgfr3^lof1/lof1^* fish and two *fgfr3^+/+^* fish (Figure 6A). After excluding outlier cells and genes, 19245 cells and an average of 1233 genes per cells were included in our analysis (Figure 6 - data 1). By applying the Seurat pipeline (Stuart et al., 2019), we identified 20 major cell clusters (Figure 6-supplement1 A). Strong cell cluster bias was not observed for our 4 samples (Figure 6-supplement1 B,G). Using a combination of known markers (Figure 6B,C, Figure 6 supplement 1 B-E and data 2 figure 6), we classified the 20 clusters into six categories: (i) cells expressing osteogenic markers (OCs; clusters 0,1,3,4,9,12, and 13), (ii) cells expressing chondrogenic markers (CHs; cluster 18), (iii) cells expressing immune markers (ICs; clusters 2, 8, 11, 16, and 19), (iv) cells expressing epidermal markers (clusters 5, 6, 10, 14, and 15), (v) cells expressing endothelial markers (cluster 17) and (vi) unannotated cells (cluster 7). Despite the lack of annotation, we noticed that nearly 10% of the cells in cluster 7 were probably pigment cells, which are numerous in the zebrafish model (Figure 6 – supplement 1F) (Saunders et al., 2019). The low observed proportion (<1%) of cells expressing classical CH marker genes (such as *col2a1, sox9, sox5* and *sox6*) was consistent with reports whereby cartilage is involved in cranial vault formation (Figure 6E) (Kanther et al., 2019).

**Figure 6:**
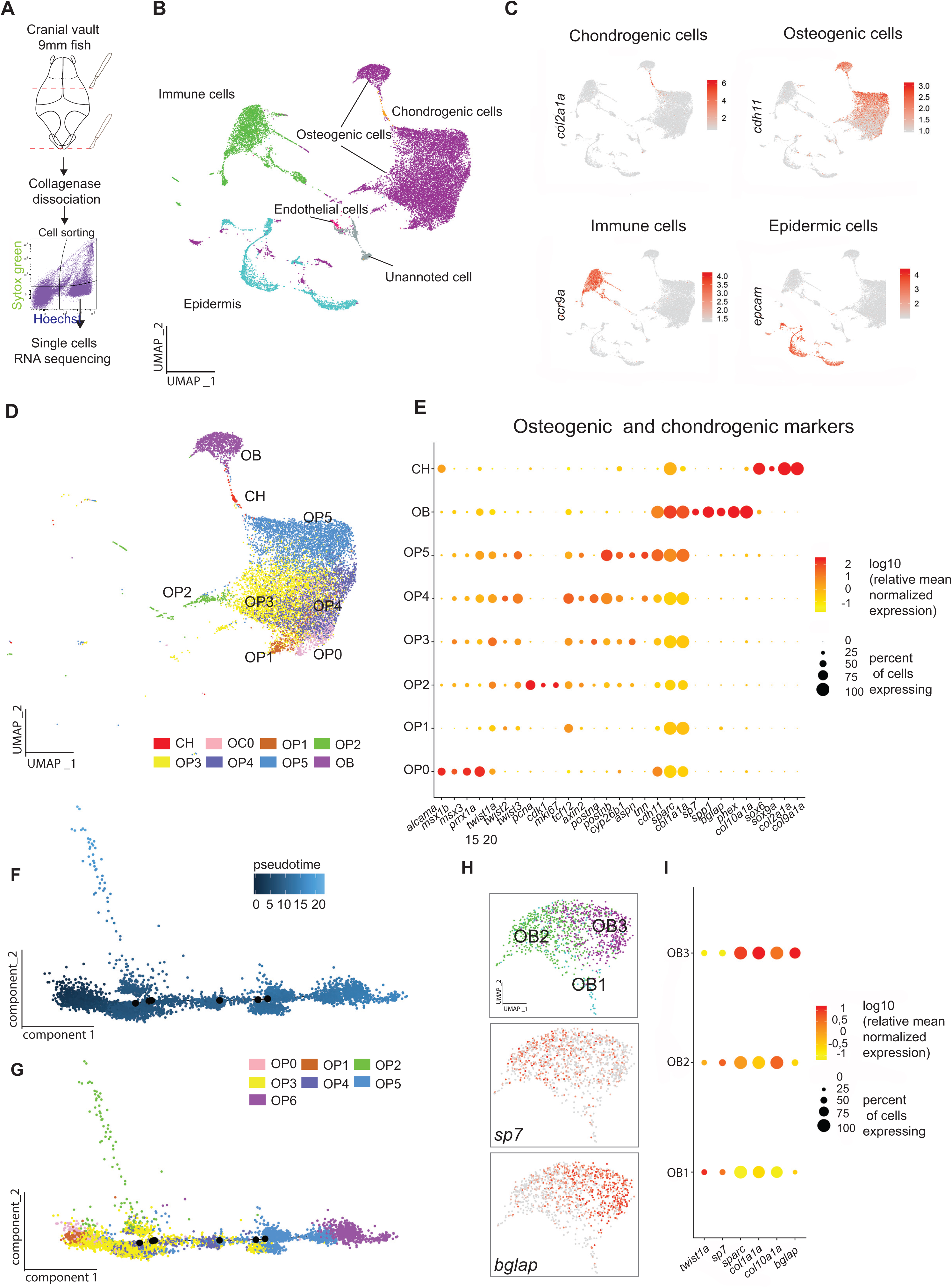
Subpopulations of osteogenic cells involving in cranial vault formation, as revealed by single-cell RNA sequencing. (A) A schematic representation of the experimental procedure. Cranial vaults were dissected, and olfactory bulb and occipital bones were removed. Cells were dissociated with collagenase and selected by sorting for SyTOX^TM^-negative and Hoechst-positive cells. (B) A UMAP analysis of pooled cells from *fgfr3^+/+^* and *fgfr3^lof1/lof1^* (n=19245), with the cell type assignments for clusters. (C) The feature plot for one known marker of each type; the color intensity corresponds to the mean normalized expression level. (D) UMAP pictures representing osteogenic (OC) and chondrogenic (CH) clusters (n=12586). (E) A dot plot showing the main marker distinguishing between OC and CH clusters. The dot diameter corresponds to the fraction of cells expressing each gene in each cluster, and the scale color corresponds to the mean normalized expression level. (F) The pseudotime trajectory of *fgfr3^+/+^* osteoprogenitor (OP) and osteoblasts (OB) cells. (G) All clusters in the trajectory are shown. (H) UMAP analysis of pooled *fgfr3^+/+^* and *fgfr3^lof1/lof1^* cells from the OB cluster (n=1166) and feature plots for *sp7* (osteoblast immature) and *bglap* (osteoblast mature) in OB clusters. (I) A dot plot showing *twist1a sp7, sparc, col1a1a, col10a1a* and *bglap* expression in OB clusters. The dot diameter corresponds to the proportion of cells expressing each gene in each cluster, as shown on the scale. The color corresponds to the mean normalized expression level.

Osteogenic cells involved in membranous ossification accounted for the largest group of cells (65%) (Figure 6 B, D and E). Cranial vault formation requires a well-orchestrated set of genes, which controls the differentiation of mesenchymal cells into mature osteoblasts; this enabled us to identify several clusters (Figure 6 D, E). Our expression analysis of the *twist* genes involved in osteoprogenitor regulation revealed that clusters 0, 1, 3, 5, 9, 12, and 13 contained osteoprogenitor cells (OP0 to OP5) (Morgan et al., 2011,Teng et al., 2018) (Figure 6E). We confirmed the presence of a large number of osteoprogenitors in the zebrafish skull at 9SL. Based on strong expression of the mesenchymal markers paired related homeobox 1a (encoded by *prrx1a)*, muscle segment homeobox 1b (*msx1b*) and muscle segment homeobox 3 (*msx3*), cluster 9 was the only cluster that contained a large number of precursor osteoprogenitor cells (OP0). The abundance of cell cycle regulators and proliferative markers in cluster 12 (Figure 6 - data 2) indicated that the latter cluster contained proliferative osteoprogenitors (OP2). Transcriptomic profiles of cluster 0 (OP3) and 3 (OP4) indicated that they both contained intermediate osteoprogenitor cells. Next, we studied the mRNA expression of *col1a1* (coding for collagen type 1a1*), cdh11* (cadherin-11) and *sparc* (osteonectin) - all of which are known to be upregulated during osteogenesis (Kii et al., 2004; Komori, 2019). We concluded that cluster 1 (OP5) contained late osteoprogenitors (Figure 6E). The increase in the expression of these genes from OP0 to OP5 reflected the progressive differentiation of osteoprogenitor cells. Osteoblasts were found in cluster 4, where we observed the expression of osteoblastic markers such as osterix *(sp7)*, osteopontin *(spp1)* and osteocalcin *(bglap)*. We were not able to classify cluster 13 (OP1).

To validate our classification of OP and OB clusters, we performed Monocle2 pseudotime trajectory analyses on *fgfr3^+/+^* cells (Trapnell et al., 2014). As shown in Figure 6G, OP and OB cells presented a relatively linear developmental progression, with few branches. The OP0 and OB clusters dominated the two ends of the progression trajectory; OB corresponded to the least mature branch in pseudotime (Figure 6). The OP5 cluster appeared to contain mostly late osteoprogenitor cells, and OP4 and OP3 contain mostly intermediate progenitors. It appeared that the cells comprising OP4 were in a less differentiated state than the cells comprising OP3. As expected, the cells forming the OP2 cluster (corresponding to proliferative osteoprogenitors) were outside the differentiation process. Interestingly, we noted that OP1 and OP0 were located at the beginning of pseudotime - nindicating that OP1 could be formed also by precursor cells. We found that the four samples contributed to all the CH, OP and OB clusters, and that there was no significant difference was observed between the genotypes (Figure 6 - supplement 1H). These results proved that absence of Fgfr3 did not affect the osteoprogenitor cell count at 9SL and confirmed our OP and OB clusters classification.

The OB cells encompassed both immature and mature osteoblasts. In order to discriminate between these two stages, we performed a new principal component analysis (PCA) and a uniform manifold approximation and projection (UMAP) analysis on cluster 4 (OB). This enabled us to identify 3 new subclusters; based on the expression of *twist1a, sp7, col1a1, col10a1a* and *bglap*, we assigned the OB1, OB2 and OB3 clusters to preosteoblasts, immature osteoblasts and mature osteoblasts, respectively (Figure 6 H,I). We noted that the proportion of mature osteoblasts (OB3) in the cranial vault was lower in *fgfr3^lof1/lof1^* fish than in *fgfr3^+/+^* fish (Figure 6 – supplemental 1 I and J). These results were consistent with the sequential live imaging data showing fewer mature osteoblasts in the cranial vault for *fgfr3^lof1/lof1^* than for *fgfr3^+/+^* at 9SL (Figure 5).

#### FGFR3 is expressed by late osteoprogenitors through to mature osteoblasts

In the wild type, *fgfr3* appeared to the *fgfr* gene most strongly expressed by osteoblasts and chondrogenic cells (Figure 7A-C); this observation was in line with the literature data (Ledwon et al., 2018; Topczewska et al., 2016). *fgfr3* was expressed in 40% of the OP5 late osteoprogenitors and in 75% of the osteoblasts. The highest expression was observed in mature osteoblasts. These data confirm the crucial role of *fgfr3* in osteoblast maturation and indicate that *fgfr3* might be also associated with earlier steps in osteogenesis. With regard to the other *fgfr* receptors, the expression of the isoform *fgfr1a* was upregulated during osteoprogenitor differentiation but *fgfr1b* was almost absent (Figure 7A). *fgfr2* was expressed weaklier than *fgfr3* during osteogenesis; as expected, it was most strongly expressed in OC0 precursor cells (Figure 7A). We noted high expression levels of *fgfr4* (whose specific role in osteogenesis is still unknown) in 47% of osteoblasts (Figure 7B). We also noted that *fgf18a* and b (specific ligands of *fgfr3*) are the most strongly expressed *fgfr* genes during cranial bone development (data not shown). Apart from the downregulation of *fgfr3*, no significant differences in the expression of three other *fgfr* genes were observed in *fgfr3^lof1/lof1^* cells (vs. the wild type). These data confirmed that *fgfr3* had indeed been knocked down in *fgfr3^lof1/lof1^* zebrafish, and showed that the other *fgfr* genes could not compensate for the absence of *fgfr3* via a putative rescue mechanism.

**Figure 7:**
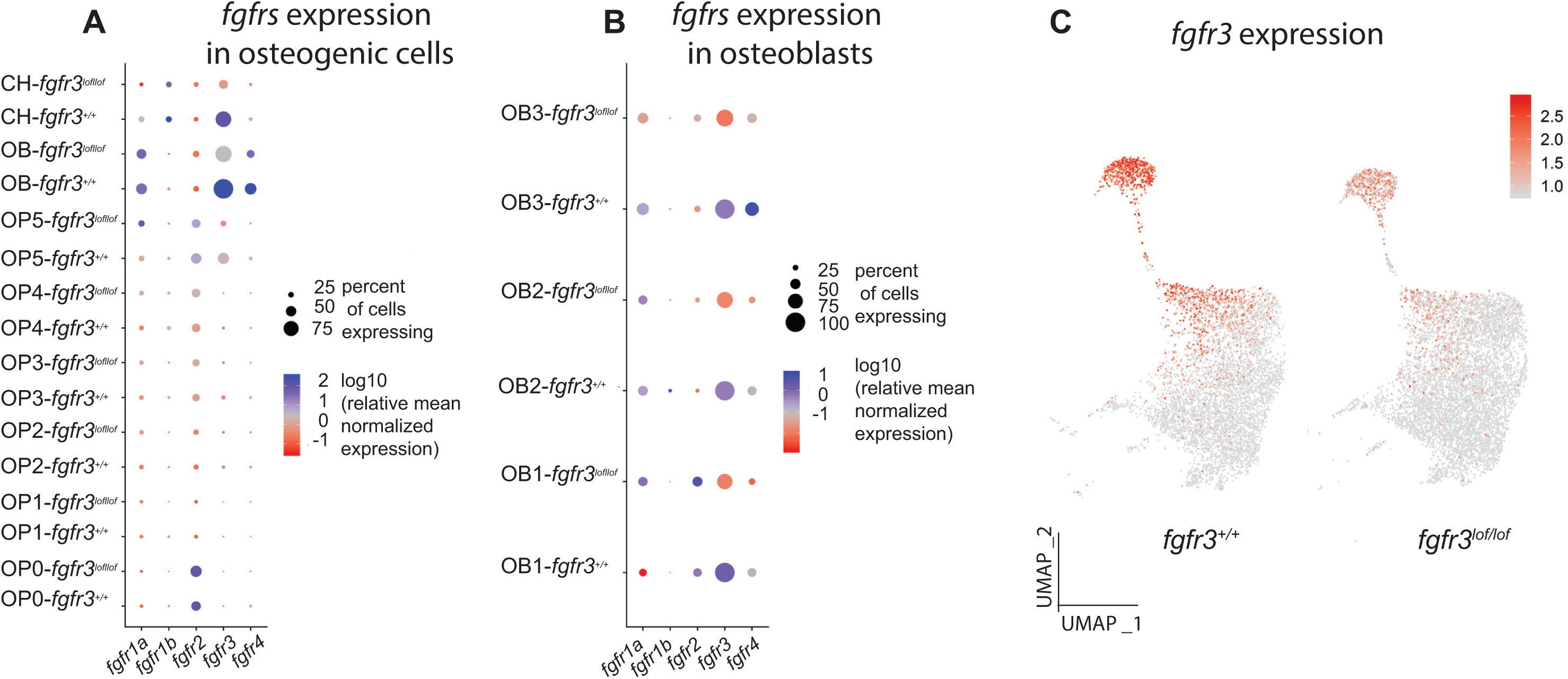
*fgfr3* is the most strongly expressed *fgfr* gene during cranial bone formation in the zebrafish. (A-B) Dot plots showing the expression of *fgfr* genes in CH, OP and OB clusters, by genotype. The dot diameter corresponds to the fraction of cells expressing each gene in each cluster, as shown on the scale. The color corresponds to the mean normalized expression level. (C) Feature plot for *fgfr3* expression in OP, OB and CH clusters. The color intensity corresponds to the mean normalized expression level.

### Fgfr3 is a positive regulator of osteoprogenitor and osteoblast differentiation

To identify changes in gene expression induced by *fgfr3* loss of function in late osteoprogenitors (OP5) and osteoblasts (OB), we performed in OP5 and OB clusters differential gene expression analysis between *fgfr3^+/+^* and *fgfr3^lof1/lof1^* in OP5 and OB clusters. To refine our list of differentially expressed genes (DEGs), we cross-checked the results obtained from each pair-wise comparison (i.e. 1-*fgfr3^+/+^* vs. 1-*fgfr3^lof1/lof1^*; 1-*fgfr3^+/+^* vs. 2-*fgfr3^lof1/lof1^*; 2-*fgfr3^+/+^* vs. 1-*fgfr3^lof1/lof1^*; 2-*fgfr3^+/+^* vs. 2-*fgfr3^lof1/lof1^*). The DEGs are listed in Figure 8 data 1.

**Figure 8:**
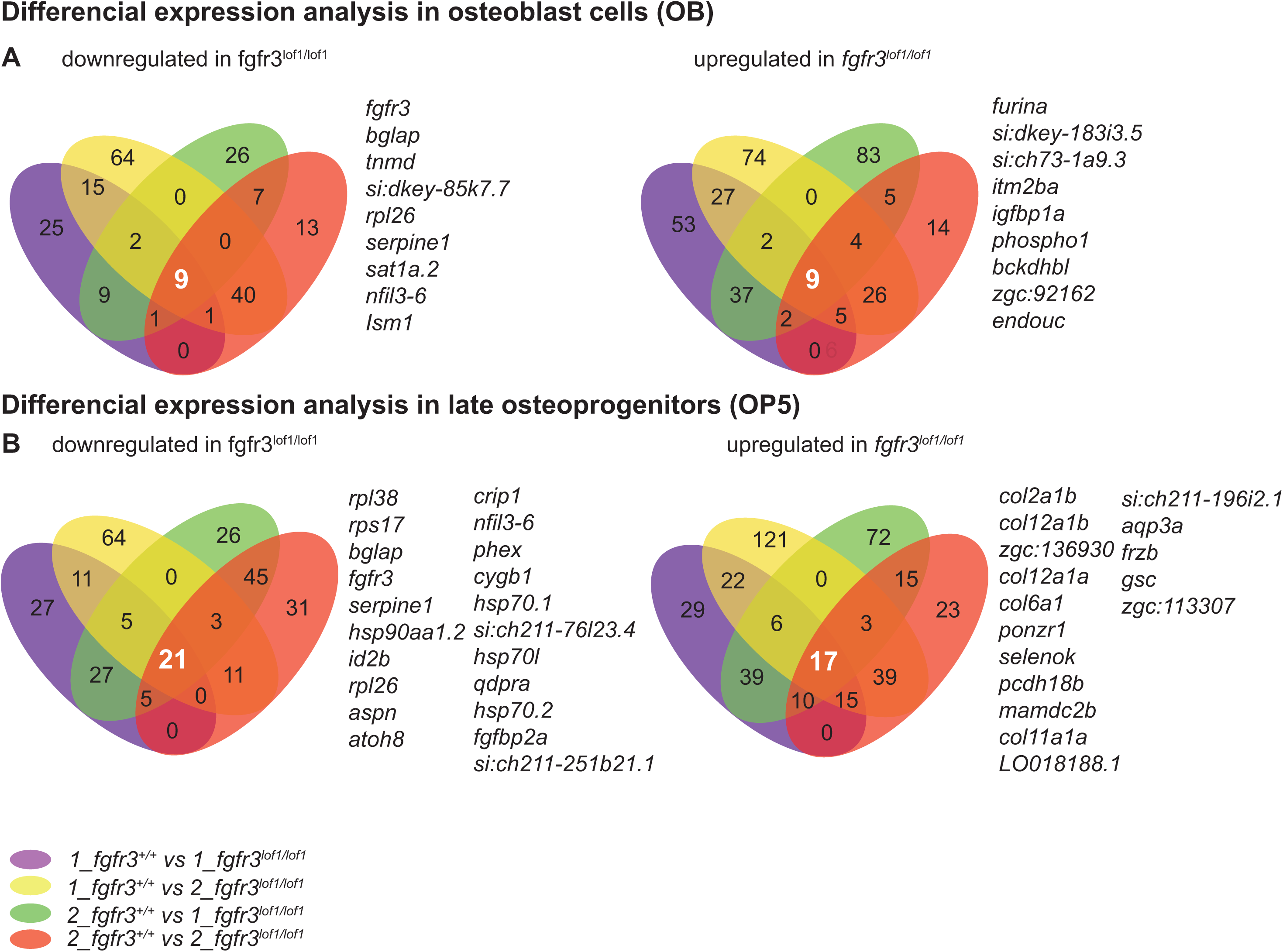
A differential gene expression analysis revealed that *fgfr3* loss of function impairs osteoprogenitor, osteoblast differentiation, and ECM constituents. (A) A Venn diagram representing the DEGs identified in the osteoblast cluster (OB). (B) A Venn diagram representing the DEGs identified in the late osteoprogenitor cluster (OP5).

As expected, we noted marked downregulation of the mature osteoblast marker *bglap* in *fgfr3 ^lof1/lof1^* cells; this confirmed the impairment in osteoblast differentiation (Figure 5) (Hauschka et al., 1989). The defective differentiation of both osteoblasts and late osteoprogenitors was supported by the dysregulation of other genes. In *fgfr3^lof1/lof1^* osteoblasts, we observed the downregulation of *sat1a.2*, coding for the spermidine/spermine N1-acetyltransferase 1a, duplicate 2 - a catabolic regulator of polyamine (Figure 8A). Polyamines and *Ssat* (the mouse ortholog of sat1a.2) are reportedly positive regulators of osteogenesis (Lee et al., 2013; Pirnes-Karhu et al., 2015). In *fgfr3 ^lof1/lof1^* late osteoprogenitors, we noted the downregulation of *inhibitor of differentiation 2* (*id2b*) coding for one of the Id proteins that promote osteoprogenitor proliferation and inhibit terminal differentiation (Peng et al., 2004). *hsp90aa1.2, hsp70.1*, and *hsp70l hsp70.2* expressions are also downregulated in *fgfr3^lof1/lof1^* late osteoprogenitors (Figure 8B). Several previous studies have shown that the mammalian orthologues *hsp70* and *hsp90* are positive regulators of osteoblast migration and differentiation (Kawabata et al., 2018; Zhang et al., 2018).

The downregulation of all these genes was in line with the defective osteoblast differentiation observed in our mutants, and highlights *fgfr3* as a positive regulator of osteoblast and osteoprogenitor differentiation.

### A lack of FGR3 affects the mineralization process and the ECM

In both late osteoprogenitors and osteoblasts, we noted that *fgfr3* loss of function impacted the expression of a large number of genes involved in mineralization and in ECM formation – both of which are fundamental for bone formation. We observed the upregulation of *phosphoethanolamine/phosphocholine phosphatase 1 (phospho1*, coding for an enzyme that helps to initiate matrix mineralization) *in fgfr3 ^lof1/lof1^* osteoblasts (Figure 8B) (Roberts et al., 2007). In *fgfr3^lof1/lof1^* late osteoprogenitors, we observed the downregulation of *asporin* (*aspn*). This small leucine-rich repeat proteoglycan/protein is expressed by osteoblast progenitor cells and promotes collagen mineralization (Kalamajski et al., 2009) (Figure 8B). *phex* is also involved in the mineralization process (Miao et al., 2001), and was downregulated in the *fgfr3 ^lof1/lof1^* OP5 cluster. Concerning the ECM, most of the genes were found in OP5. Indeed, we observed the upregulation of several genes coding for collagen proteins (*col2a1b, col12a1b, col12a1a* and *col6a1*) (Figure 8B). We also noted the upregulation of *fibromodulin (zgc:113307;fmod)*, a collagen-binding small leucine-rich proteoglycan regulating collagen fibril formation in the *fgfr3^lof1/lof1^* OP5 population (Figure 8B). We observed the downregulation of *insulin-like growth factor binding protein 1a (igfbp1a*) in osteoblasts (Figure 8A). Interestingly, insulin-like growth factor 1 signaling is dysregulated in single-suture craniosynostosis and control ECM focal adhesion and the human homolog *igfbp1a* is known to be expressed in osteoblastic cells, and to inhibit collagen gene expression by blocking the effect of insulin growth factor 1 (Stamper et al., 2011; Rosen et al., 1994).

Our analysis of the DEGs confirmed our starting hypothesis, i.e. that *fgfr3* loss of function perturbed the differentiation of both late osteoprogenitors and immature osteoblasts. These defects are associated with marked modifications of the ECM and abnormal mineralization, as evidenced by Alizarin red S staining. Impairment of late osteoprogenitor differentiation and ECM synthesis might explain the defect in immature expansion observed in *fgfr3 ^lof1/lof1^* zebrafish.

## Discussion

By leveraging our new zebrafish *fgfr3^lof/lof^* mutant, we assessed the involvement of Fgfr3 in cranial vault development and membranous ossification. *fgfr3* loss of function induced marked anomalies in the craniofacial skeleton, with microcephaly, Wormian bone formation, and small frontal and parietal bones. As *Fgfr3* knock-out mice do not present obvious craniofacial anomalies, the *fgfr3^lof/lof^* zebrafish appears to be the first animal model to display much the same craniofacial defects (microcephaly and Wormian bones) as in human *FGFR3* loss-of-function related diseases (Deng et al., 1996; Eswarakumar and Schlessinger, 2007; Makrythanasis et al., 2014; Toydemir et al., 2006).

The different craniofacial phenotypes observed in humans, zebrafish and mice with *FGFR3* loss-of-function mutations might be due to interspecies differences in the *fgfr3* gene’s expression profile. In humans, *FGFR3* is expressed during cranial vault formation and suture formation. On the fertile side of the sutures, *FGFR3* is expressed less strongly than *FGFR2* and *FGFR1* (Delezoide et al., 1998). In the zebrafish, previous studies and our present single-cell transcriptomic analysis revealed that expression levels of *fgfr3* are higher than those of *fgfr1a, fgfr1b* and *fgfr2* in (i) late osteoprogenitors and osteoblasts during cranial vault development and (ii) the osteogenic cells lining the suture’s mineralization front (Topczewska et al., 2016). The situation is different in the rodent; *Fgfr3c* is almost completely absent from developing cranial bones, and is more weakly expressed than *Fgfr2c* and *Fgfr1c* in osteogenic fronts and sutures (Molténi et al., 1999; Rice et al., 2003). *Fgfr2* and *Fgfr1* are known to be regulators of cranial bone formation, whereas *Fgfr3*’s role is not well understood (Ornitz and Marie, 2019). Earlier studies evidenced the upregulation of *Fgfr1c* and *Fgfr2c* expression in osteoblastic cells isolated from murine *Fgfr3^-/-^* bone marrow stromal cells (Valverde-Franco et al., 2004). These data indicate that *Fgfr1c* and *Fgfr2c* can compensate for the absence of *Fgfr3c* during cranial vault formation in the mouse. In our single-cell analyses, expression levels of *fgfr1a*, *fgfr1b* and *fgfr2* were similar in *fgfr3^lof1/lof1^* zebrafish and controls. These findings demonstrated that the absence of *fgfr3* was not compensated for in *fgfr3^lo1/lof1^* fish.

Cranial vault formation starts with the formation of ossification centers via the condensation of mesenchymal cells and the latter’s differentiation into osteoprogenitors, immature osteoblasts and then mature osteoblasts. Bone growth combines the production of osteoids, their mineralization by osteoblasts, and the continued differentiation of mesenchymal cells at the periphery. The role of Fgfr3 during osteoblast differentiation is subject to debate; a recent review suggested that Fgfr3 (i) indirectly regulates osteoblasts during cranial vault development, and (ii) has paracrine properties in the context of endochondral ossification (Ornitz and Marie, 2019; Matsushita et al., 2009; Mugniery et al., 2012; Wen et al., 2016). Other *in vitro* studies have demonstrated a direct effect of *fgfr3* on osteoblast differentiation (Su et al., 2010; Valverde-Franco et al., 2004). Nevertheless, the absence of Fgfr3 and the expression of a constitutively activated form of Fgfr3 in mouse cells lead to premature osteoblast differentiation and impairment of the mineralization process. Here, we took advantage of the zebrafish model to perform live *in vivo* imaging. We found that Fgfr3 regulates the expansion and differentiation of immature osteoblasts during membranous ossification. Our results evidenced high expression levels of *fgfr3* in osteoblasts and (to a lesser extent) late osteoprogenitor cells - indicating that Fgfr3 acts directly on osteogenesis in the zebrafish.

The key features of cranial vault bone formation include the relative locations of ossification centers of the individual vault bones. In our study, we observed that the primary ossification center in both mutant and control was indeed initiated in the early stages of development. We showed that the impairment of ossification center expansion is consistent with the increase in cell cycle duration observed in mutant immature osteoblasts at 9SL, and leads to a decrease over time in the immature osteoblast count. One can reasonably assume that this defect in immature osteoblast expansion results from a defect of new osteoblast recruitment, which depends on the osteoprogenitor population. It is important to note that the maturation of osteoblasts from 9SL to 11SL was also blocked, as revealed by the presence of a huge number of immature osteoblasts. The impairment of each step in osteogenesis only appeared once ossification growth had been initiated - suggesting that the cells’ position with regard to the ossification center is crucial, and that cell-cell communication was defective. Consistently, our differential gene expression analysis revealed that the absence of functional Fgfr3 is associated with changes in the production of ECM constituents by osteoblasts and osteoprogenitors. The ECM proteins have an important role in cell-cell communication, including matrix bridge formation and morphogen regulation. It has been well established that the osteoblasts in the ossification center secrete morphogens that diffuse into neighboring cells and activate mesenchymal cell differentiation (Garzón-Alvarado et al., 2013; Lee et al., 2015). Here, the DEGs included *fmod* and *aspn*, both of which code for small leucine-reach repeat proteoglycans (fibromodulin and asporin, respectively) expressed in the ECM of bone, cartilage and tendons. It is known that in osteoarthritic cartilage, fibromodulin and asporin are able to modulate TGF-beta signaling by sequestrating TGFb1 within the ECM (Embree et al., 2010; Kalamajski et al., 2009). Fmod also controls the fate of tendon progenitor cells by modulating bone morphogenic protein activities (Bi et al., 2007). In the present context, we hypothesized that dysregulation of *fmod* and *aspn* impaired the shaping of the morphogenetic fields required for proper bone development. Moreover, we observed the upregulation of *col6a1, col12a1a* and *col12a1b*; these collagen gene products are known to be involved in the formation of matrix bridges between cells during osteogenesis, and are essential for proper bone formation (Izu et al., 2016, 2011). On the basis of these data, we assume that the changes in ECM components in growing bone caused by the absence of *fgfr3* impair cell-cell communication and thus disturb the recruitment of new osteoblasts. Further in-depth analysis of the ECM and signaling pathways in fgfr3*^lof/lof^* fish will be necessary to confirm our hypothesis.

Interestingly, the absence of Fgfr3 in zebrafish (as in humans) leads to Wormian bone formation (Toydemir et al., 2006). In humans, these bones are found in the general population but are primarily associated with several genetic bone disorders involving skull deformation (hydrocephaly, craniosynostosis, cleidocranial dysplasia, osteogenesis imperfecta, etc.) (Bellary et al., 2013; Sanchez-Lara et al., 2007). Our observations revealed that in contrast to the zebrafish *sp7* knock-out model showing that Wormian bone formation resulted from the increase in early osteoblast recruitment, in *fgfr3^lof/lof^* fish the ectopic bones appeared as a result of the bone formation delay (Kague et al., 2016). Our data highlighted the complexity of Wormian bone formation, a topic that warrants further investigation.

In conclusion, the *fgfr3^lof1/lof1^* zebrafish is the first animal model to have enabled an *in vivo* analysis of Fgfr3’s role in intramembranous ossification. Thanks to the late development and accessibility of the cranial vault, the *fgfr3^lof1/lof1^* model provides unprecedented access to the mechanism of bone formation in a live animal. Moreover, the phenotype displayed by the mutant fish is closer to that seen in humans than that observed in the mouse (e.g. the presence of Wormian bones and microcephaly). Our integrated analyses in the zebrafish (including *in vivo* experiments and single-cell studies) clearly demonstrated that *fgfr3* is a positive regulator of osteogenesis during membranous ossification. The results of our transcriptomic single-cell analysis demonstrated for the first time that Fgfr3 is able to regulate the ECM during skull vault formation.

We expect these data to be of value in understanding skull vault anomalies observed in patients with FGFR3-related disorders.

## Materials and Methods

### Zebrafish husbandry and transgenic strains

Zebrafish were raised, hatched and maintained in an animal house at 28°C (approval number 2018080216094268), in compliance with the EU Directive 2010/63/EU for animals. Fish were studied at various developmental stages, and the standard length (SL) was measured from the snout to the caudal peduncle (Parichy et al., 2009). The *Tg(sp7:mCherry)* and *Tg(bglap:GFP)* fish lines have been previously described; they labeled immature osteoblasts and mature osteoblasts, respectively (Knopf et al., 2011). The *Tg(col2:mCherry)* fish line was used to label chondrocytes (Hammond and Schulte-Merker, 2009).

### BGJ398 treatment

*Tg*: (*sp7: mCherry*) zebrafish with an SL of 7mm at treatment initiation received intraperitoneal administrations of BGJ398 (2 mg/kg body weight) (infigratinib; LC Laboratories) or vehicle (3.5 mM HCl, DMSO 3%) every other day for 15 days. After treatment, fish were euthanized by balneation in 75 µg/L benzocaine solution. Photographs were taken with an M2015FCA stereomicroscope equipped with a Leica DFC300G camera and a filter for mCherry. The SL and fontanelle area were measured using Fiji software (Schindelin et al., 2012).

### Generation of the CRISPR/Cas9 Fgfr3^lof^ zebrafish line

Two loss-of-function *fgfr3* lines were established by using clustered regularly interspaced short palindromic repeats (CRISPR)/CRISPR-associated protein 9 (Cas9) genome editing. Single guide RNAs were designed using the CRISPOR online tool (http://crispor.tefor.net). The *fgfr3* targets in this study were 5′-TTTGTGCTCTCTGTGTCAGCGG-3′ and 5′-GCCCACCACTACCGTCTGAT-3’ for the *fgfr3*^lof1^ and *fgfr3*^lof2^ lines, respectively.

Injection mixtures included 12 µM of Cas9 protein and 200 pg of guide RNA. Both lines present a premature stop codon upstream of the transmembrane domain, leading to the loss of the tyrosine kinase domain. The *fgfr3*^lof1^ line carries a 3-base-pair deletion (2 bases in intron 8-9 and 1 base in exon 9), leading to a frameshift after amino acid 372 and a premature stop at position 377. The *fgfr3*^lof2^ line carries a 4-base-pair deletion, leading to a frameshift after amino acid 277 and a premature stop at position 278. We crossed *fgfr3* ^+/lof^ females and *fgfr3^+flof^* females to obtain *fgfr3 ^lof/lof^*, and *fgfr3* ^+^*^f+^* siblings as controls for phenotypic analyses.

### Genotyping

Genomic DNA was isolated from caudal fin biopsies. Tissues were lysed in 10mM Tris– HCl, pH 8.5, 2mM EDTA pH8, 0.2% Triton X-100 and 250 µg/ml proteinase K (Sigma Aldrich) for at least 1h at 55°C. The samples were genotyped in heteroduplex mobility assay using the following primers *fgfr3*^lof1^_F: 5’-TGAACGATCCCAGGAATCCC-3’ and *fgfr3*^lof1^_R: 5’-GGTGAGAATGAAGAGCACGC-3’, *fgfr3*^lof2^_F: 5’-CAACTCGATGTGCTGGGTAT-3’ and *fgfr3*^lof2^_R: 5’-GCTGAGCATCACTGTACACC-3’.

### Skeletal staining

#### Alizarin red S staining of 9dpf larvae

Bones were stained with Alizarin red S (Sigma Aldrich), using the method described by Walker and Kimmel (Walker and Kimmel, 2007).

#### Alizarin red S staining of larvae (7SL to 11SL)

zebrafish were sacrificed and fixed overnight in 96% ethanol. Soft tissues were dissolved by placing fish in 2%KOH v/v for 24 hours. Next, the bones were stained overnight in a 0.075% Alizarin red S w/v 1% KOH v/v solution. Lastly, to remove pigment and remaining tissue, the fish were placed in 20% glycerol v/v and 1% KOH for at least a week.

#### Alcian blue and Alizarin red S staining of 3-month-old fish

Zebrafish were sacrificed and fixed overnight in 96% ethanol. Fish were stained overnight at room temperature (RT) with a solution of 0.1% Alcian blue 8GX (Sigma Aldrich) w/v, 70% ethanol v/v, and 20% glacial acid acetic v/v. Fish were rinsed overnight in 96% ethanol, and soft tissue was dissolved by placing fish in 2%KOH v/v for 72 hours. The bones were then stained overnight in a 0.075% Alizarin red S w/v, 1% KOH v/v solution. Scales, fat and organs were dissected manually. Lastly, the fish were placed in 20% glycerol v/v, 1% KOH for several weeks to remove the remaining tissue.

#### Vital staining with calcein green and Alizarin red

Zebrafish at 9 SL were used for live staining. Using water from the fish facility, 0.003% Alizarin red S and 0.005% calcein green (Sigma Aldrich) stains were applied sequentially in overnight incubations. The Alizarin red S stain was applied at 9SL, and was followed by calcein green staining 7 days later.

The fish were imaged with an M2015FCA stereomicroscope equipped with both a Leica DFC300G camera (a mCherry filter was used to image Alizarin Red, and a GFP filter was used to image calcein) and a Leica MC 170HD camera for bright-field images.

### X-ray and µCT analyses

The fish’s whole skeletons were X-rayed using a Faxitron (MX-20DC12). Vertebrae were measured on the radiographs, using Image J software. Micro computed tomography (voxel size: 2 μm) on a Skyscan 1272 system (Bruker)) was used to evaluate the skull morphology in *fgfr3^lof1/lof1^* and *fgfr3^+/+^* fish (n=5 per group). The skull of each fish was dissected, fixed in 4% phosphate-buffered formaldehyde for at least 24 hours, dried in a series of increasingly concentrated ethanol solutions and, lastly, dried in air. To reduce movement, specimens were glued to a sample holder during the µCT scan. Scans were acquired at 40 kV and 240 μA with a 0.25 mm aluminum filter. Ring artifacts and beam hardening corrections were the same for all scans, during reconstruction with NRecon software (Bruker) according to previously established protocols(Busse et al., 2019; Fiedler et al., 2018; Suniaga et al., 2018).

### Histological analyses and *in situ hybridization*

Whole fish were fixed overnight in 4% paraformaldehyde at 4°C, decalcified in a 0.5 M EDTA pH 8.0 solution for 3 weeks, and embedded in paraffin. Serial sections of 6 mm were stained with hematoxylin–eosin or underwent an *in situ* hybridization analysis. *col1a1* digoxigenin (DIG) riboprobes were obtained with following primers: F-TGAGGCCTCCCAGAACATTA and R-TTGGTCAACACTGGGAATCG. *In situ* hybridization was performed on sections deparaffinated with Histo-Clear^TM^ solution (National Diagnostics), rehydrated in a series of ethanol/PBS solutions (Gibco), permeabilized with proteinase K (25 μg/ml), prehybridized, and then hybridized overnight at 65°C in hybridization mixture (50% formamide, 5× standard saline citrate [SSC], 0.1% Tween 20, 100 μg/ml heparin, 100 μg/ml tRNA in water). After a series of washes in 50% SSC/formamide and 0.2%SSC/PBS with 0.1% Tween (PBST), sections were incubated in blocking solution (0.2% Tween 20, 0.2% Triton X-100, and 2% sheep serum in PBST) and incubated overnight at 4°C with AP-conjugated anti-DIG antibodies diluted 1:4000 in blocking solution. Sections were washed in PBST, soaked in staining buffer (100 mM NaCl, 100 mM Tris-HCl, pH 9.5, 0.1% Tween 20 in water), and then incubated in nitroblue tetrazolium/5-bromo-4-chloro-3-indolyl phosphate solution (Roche).

### Immunohistochemistry and EdU Labeling

For 5-ethynyl-2′-deoxyuridine (EdU) incorporation experiments, 9SL fish were intraperitoneally injected with 1 µl of 10 mM EdU solution per 15 mg using a microinjector. One hour later, the fish were then killed by over-anesthesia. Next, the calvaria were dissected out and processed as described below for immunohistochemical assessment. EdU was detected with the EdU Click-iT Plus EdU Alexa Fluor 488 Imaging Kit (Life Technologies), according to the manufacturer’s instructions.

For the immunohistochemical assessment, dissected calvaria from 9SL or 11SL fish were labeled with antibodies against mCherry (Takara, 1/100), GFP (Roche, 1/100) and pH3 (Abcam; 1/100). The calvaria were fixed in 4% paraformaldehyde (Electron Microscopy Sciences) in PBST, depigmented using Bloch *et al.’s* protocol (Bloch et al., 2019), and permeabilized for 5 minutes in 10 µg/ml proteinase K solution w/v in 0.5% Triton X-100 PBS (PBTr). The samples were rinsed 3 times in PBTr and post-fixed in 4% PFA for 20 minutes at room temperature. After 3 washes in PBTr, the samples were incubated at room temperature for at least 1 h in a blocking solution containing 10% normal goat serum, 1% DMSO, and 0.5% Triton X-100 in PBS. Next, samples were incubated with primary antibody diluted in blocking solution 4°C for 3 days with gentle shaking. Lastly, primary antibodies were detected using appropriate secondary antibodies coupled to Alexa-Fluor 488 and Alexa Fluor 594 (1:100; Molecular Probes) and diluted in PBTr for 2 days at 4°C.

After a permeabilization step, Terminal deoxynucleotidyl transferase dUTP nick end labeling (TUNEL) TUNEL assays were performed with the Deadend Fluorometric TUNEL system (Promega), according to manufacturer’s instructions. A Zeiss Z1 light sheet microscope was used to image *in toto* the calvaria processed for immunohistochemical assessment, TUNEL assays, and EdU experiments. The co-localization analysis of parietal bones was performed using Imaris software (version 9.1.2). The mCherry-positive volume was used to extract the volumes of interest for analysis.

### Sequential imaging

For sequential live imaging, fish were anesthetized with benzocaine (75 µg/L) and mounted in a Petri dish in which a well had been carved out of 1% low-melting-point agarose. Sequential pictures were taken in the same orientation, using a Yokogawa CSU-X1 spinning disk scanner coupled to a Zeiss Observer Z1 inverted microscope and a DP72 camera controlled by Zen Blue software (Olympus).

### Quantitative real-time PCR

Total RNA from 6-week-old fish was extracted with Trizol reagent (Thermo Fisher Scientific), according to the manufacturer’s protocol. First-strand complementary DNA (cDNA) was synthesized from 1 µg of RNA using the SuperScript First-Strand Synthesis System (Thermo Fisher Scientific), according to the manufacturer’s instructions. Quantitative PCR was performed with PowerUp SYBR Green Master Mix (A25777, Thermo Fisher Scientific). The following primers were used: m*βactin* (F 5’-GCCAACAGGGAAAAGATGAC-3’; R 5’-GACACCATCACCAGAGTCCA-3’), *fgfr1a* (F 5’-GATGCCACGGAGAAAGACCT-3’ and R 5’-AGCAGCAAACTCCACGATGA-3’), *fgfr1b* (F 5’-GGTCTCCTGCGCATACCA-3’ and R 5’-CAGGCCGAAGTCTGCTATCT-3’), *fgfr2* (F 5’-AAAGAGGGTCATCGCATGGA-3’ and R 5’-ATCCTCCACCAACTGCTTGA-3’) and *fgfr3* (F 5’-GGCACCAGAAGCACTGTT-3’ and R 5’-CACTGGGATACCTGGATACG-3’). In the qPCR, expression of *fgfr1a*, *fgfr1b*, *fgfr2, and fgfr3* were normalized against *βactin*, gene expression was quantified using the equation 2^−ΔΔCT^, and the fold-change in mRNA expression was calculated relative to the mRNA level in one of the controls (set to 1).

### Cell dissociation, fluorescence-activated cell sorting, single-cell sample preparation, and RNA sequencing

The cranial vault was isolated from the head of 9SL zebrafish and dissected in PBS. The olfactory bulbs, the anterior parts of the frontal bones, and the suboccipital bones were removed (Figure 6 A). The soft tissue was partially degraded by incubation with 0.2% collagenase (Sigma-Aldrich-C5138) and 4 mM EDTA in PBS (Gibco) for 10 min at 28°C. After a wash in PBS, cells were dissociated for 40 min at 28°C in 0.2% collagenase in PBS. The solution was strained using a 40 μm cell strainer (Corning), and the cells were washed in α-MEM supplemented with 10% FBS. Prior to flow cytometry, the cells were suspended in FacsMax (amsbio) and stained with Sytox^TM^ green nucleic acid stain (1:100000, Invitrogen) and Hoechst (1:5000, Thermo Fisher). To sort the cells, beads (Sony-LE-B3001) were used to select events larger than 3 µm. Hoechst-positive cells (cells with a nucleus) and Syber-green-negative cells (live cells) were sorted on an Aria 2 sorp cell sorter (Beckson Dickinson).

The scRNA-seq libraries were generated with a Chromium Single Cell 3′ Library & Gel Bead Kit v.3 (10x Genomics), according to the manufacturer’s instructions. Briefly, cells were counted, diluted at 1000 cells/µL in PBS+0.04% BSA, and 6500 cells were loaded in the 10x Chromium Controller to generate single-cell gel beads in an emulsion. After reverse transcription, the gel beads in the emulsion were disrupted. Barcoded complementary DNA was isolated and amplified in PCRs. Following fragmentation, end repair and A-tailing, sample indexes were added during index PCR. The purified libraries were sequenced on a Novaseq 6000 (Illumina) on paired end strands (read 1: 28bp; read 2: 91bp), with a mean read depth of 25000 reads per cell.

Fastq files from the scRNA 10X libraries were processed using the CellRanger Count pipeline with its default parameters. Reads were aligned against the GRCz11-3.0.0 reference transcriptome. The RNA data quality control and downstream analysis were performed using the Seurat R package (version 3.0.2) and the standard Seurat v3 integration workflow (Stuart et al., 2019). We filtered cells that had (i) unique feature counts over 2000 or below 500, or (ii) more than 20% of mitochondrial counts. After data filtering, we obtained an expression matrix with 21176 genes and 19245 cells. The matrix’s dimensions were reduced by running the 20 significant principal components against scaled data. To identify cell populations, we started with a clustering resolution of 0.6, used the expression of a panel of gene markers to annotate clusters, and then decreased the resolution to 0.2 to refine the clusters.

Data were visualized using the RunUMAP function that computed a new matrix of two-dimensional coordinates for each cell, based on the distance between clusters. Next, we performed a new PCA and a UMAP analysis of cluster 4 (OB), corresponding to osteoblasts. A clustering resolution of 0.2 was used to discriminate between immature and mature osteoblasts.

We used the FindMarkers function to compute differentially expressed genes (DEGs) for each of the identity classes in the dataset. Differential expression was defined using a parametric Wilcoxon rank sum test and a significance level of p<0.05. To refine our lists of DEGs and reduce the background noise, we cross-checked the results obtained from each pair-wise comparison (1-*fgfr3^+/+^* vs. 1-*fgfr3^lof1/lof1^*; 1-*fgfr3^+/+^* vs. 2-*fgfr3^lof1/lof1^*; 2-*fgfr3^+/+^* vs. 1-*fgfr3^lof1/lof1^*; 2-*fgfr3^+/+^* vs. 2-*fgfr3^lof1/lof1^*). Venn diagrams were drawn using Venny (https://bioinfogp.cnb.csic.es/tools/venny/index.htm).

### Pseudotime analysis

The top 1000 highly dispersed genes among the osteogenic lineage cell clusters (OP0, OP1, OP2, OP3, OP4, OP5, OB) in WT fish were chosen as feature genes to resolve pseudotemporal trajectories using the setOrderingFilter, reduceDimension, and orderCells functions of Monocle (v2.8.0). We used the default parameters (except for max_components = 4 and norm_method = log) to generate the 3D trajectory during dimensionality reduction.

### Statistical analysis

All statistical analyses were performed using Prism 6 (GraphPad Software), with the statistical tests and corresponding two-tailed P-values reported in the figure legends: *, p<0.05, **, p<0.01, ***, p<0.001 and ****, p<0.0001 were considered to be statistically significant. All values are quoted as the mean +SEM.

## Acknowledgements

We are grateful to Dr. Chrissy Hammond for providing the Tg(*col2:mCherry*) fish line and Dr.Gilbert Weidinger for providing the *Tg(sp7:mCherry)* and *Tg(bglap:GFP)* fish lines. We thank all the members of the Cell Imaging facility at Necker Children’s Hospital for technical assistance with confocal microscopy and for helpful discussions. We thank the staff at the Institut Curie’s next-generation sequencing facility for the high-throughput sequencing, Camille de Cevins of Menager lab, and staff at the Institut Imagine’s animal facility. We also thank the members of the Legeai-Mallet lab for helpful discussions. With regard to funding, the high-throughput sequencing was funded by grants ANR-10-EQPX-03 and ANR10-INBS-09-08 from the *Agence Nationale de le Recherche* (*Investissements d’Avenir* program) and a grant from the Canceropole Ile-de-France to the next-generation sequencing facility at the Institut Curie. Some of the work presented here was funded by the European Community’s Seventh Framework Programme under grant agreement 602300 (the SYBIL program-https://www.sybil-fp7.eu/- is funded by the MRC (MC_UU_000007/9), the European Research Council (ZF-MEL-CHEMBIO-648489) and the L’Oreal – Melanoma Research Alliance (401181). Emilie Dambroise received postdoctoral fellowship from *l’Association des Gueules Cassées*.

## Competing interests

None declared.

