## Supplementary material for "FGFR3 is a positive regulator of osteoblast expansion and differentiation during zebrafish skull vault development": Figures supplement

### Legends:

**Figure 1 supplement 1: Fgfr3's amino acid sequence is highly conserved between zebrafish and humans.**

Human FGFR3 and zebrafish Fgfr3 protein sequences were aligned using CLC software. Identical residues are depicted in blue. The extracellular domain is shaded in yellow. The transmembrane domain (from position 399 to 418) is shaded in green. The intracellular domain is shaded in purple, and encompasses the tyrosine kinase domain (in dark purple).

**Figure 2 supplement 1: Three-month-old *fgfr3$^{lof2/lof2}$ fish* present craniofacial and axial skeletal anomalies**

(A) Schematic representation of FGFR3 receptor, with the position of the lof2 mutation. IG: Immunoglobulin domain; TM: transmembrane domain; TK: tyrosine kinase domain. Standard length (*fgfr3*$^{+/+}$ n=9; *fgfr3*$^{lof2/lof2}$ n=10) (C) Images of sibling *fgfr3*$^{+/+}$ and *fgfr3*$^{lof2/lof2}$ fish at 3 months of age, demonstrating that *fgfr3*$^{lof2/lof2}$ fish present flat face, microcephaly, and a low SL. (D) X-rays of *fgfr3*$^{+/+}$ and *fgfr3*$^{lof2/lof2}$ fish show severe anomalies of the craniofacial skeleton and swim bladder in the mutant. (D) A dorsal view of the cranial vault stained with Alizarin red S, highlighting defects in the frontal and parietal bones and the presence of ectopic bones (white stars) in the mutant. BF: bright field; A: anterior; P: posterior; D: dorsal; V: ventral; f: frontal; p: parietal; ns: non-significant. P-values were determined using Student's t-test; the error bars correspond to the standard error of the mean.

**Figure 2 supplement 2: Fgfr3 loss of function did not affect the appendicular skeleton**

(A) Alcian blue and Alizarin red S staining of dorsal, anal, pelvic and pectoral fins of 3 month-old *fgfr3*$^{+/+}$ and *fgfr3*$^{lof1/lof1}$ fish. (B-E) Length of the dorsal, anal, pelvic and pectoral fins (*fgfr3*$^{+/+}$ n=6; *fgfr3*$^{lof1/lof1}$ n=6) ns: non-significant. P-values were determined using Student's t-test; the error bars correspond to the standard error of the mean.

**Figure 3 supplement 1: Analyses of the skeletal phenotype in 6–month-old *fgfr3$^{lof1/lof1}$* fish**

(A) Image of 6-month-old sibling *fgfr3$^{+/+}$* and *fgfr3$^{lof1/lof1}$* fish, showing that *fgfr3$^{lof1/lof1}$* fish present a flat face and microcephaly. (B) X-rays of *fgfr3$^{+/+}$* and *fgfr3$^{lof1/lof1}$* fish, showing severe anomalies of the craniofacial skeleton and the swim bladder in the mutant. (C) A dorsal view of cranial vault stained with Alizarin red S, highlighting frontal and parietal bone defects and the presence of ectopic bones (white stars) in mutants (D) Sections stained with hematoxylin-eosin reagent show the absence of a sagittal suture in *fgfr3$^{lof1/lof1}$* fish. (E) Sections stained with hematoxylin-eosin reagent show the absence of a metopic suture in *fgfr3$^{lof1/lof1}$* fish. A: anterior; P: posterior; D: dorsal; V: ventral; f: frontal; p: parietal.

**Figure 3 supplement 2: Analyses of the larval *fgfr3$^{lof1/lof1}$* viscerocranium**

(A) Alizarin red S staining of *fgfr3$^{lof1/lof1}$* fish and their siblings at 9dpf. The *fgfr3$^{lof1/lof1}$* fish did not present delayed mineralization of the branchial arch. (B and C) There were no differences in head width or length (*fgfr3$^{+/+}$*: n=13; *fgfr3$^{lof1/lof1}$*: n=13) (D) *col2*: *mCherry* expression in chondrocytes in 14 dpf *fgfr3$^{lof1/lof1}$* and sibling larvae. (E-G) No There were no differences in the length of the Meckel, palatoquadrate and ceratohyal cartilages (*fgfr3$^{+/+}$* n=9; *fgfr3$^{lof1/lof1}$* n=9). ns: non-significant; A: anterior; P: posterior; D: dorsal; V: ventral; M: meckel; Pq: Palatoquadrate; Ch: ceratohyalate. P-values were determined using Student's t-test; the error bars correspond to the standard error of the mean.

**Figure 6 supplement 1: Subpopulations of osteogenic cells involving in cranial vault formation, as revealed by single-cell RNA sequencing**

(A) UMAP analysis of pooled cells from *fgfr3$^{+/+}$* and *fgfr3$^{lof1/lof1}$* (n=19245) (B) A UMAP analysis of each sample 1-*fgfr3$^{+/+}$* (n=3651), 2-*fgfr3$^{+/+}$* (n=5828), 1-*fgfr3$^{lof1/lof1}$* (n=6623) and 2-*fgfr3$^{lof1/lof1}$* (n=3143). (C-F). Dot plots showing the main markers distinguishing between immune, epidermal, endothelial and pigment cells in all clusters. The dot diameter corresponds to the fraction of cells expressing each gene in each cluster, as shown on the scale. The color corresponds to the mean normalized expression level. (G) The proportion of cells in each category, by genotype. (H) The proportion of cells in each osteogenic and chondrogenic cluster, by genotype. (I) A UMAP analysis of pooled cells from the OB cluster in *fgfr3$^{+/+}$* (n=605) or *fgfr3$^{lof1/lof1}$* fish (n=561) (J) The proportion of cells in each OB cluster, by genotype.

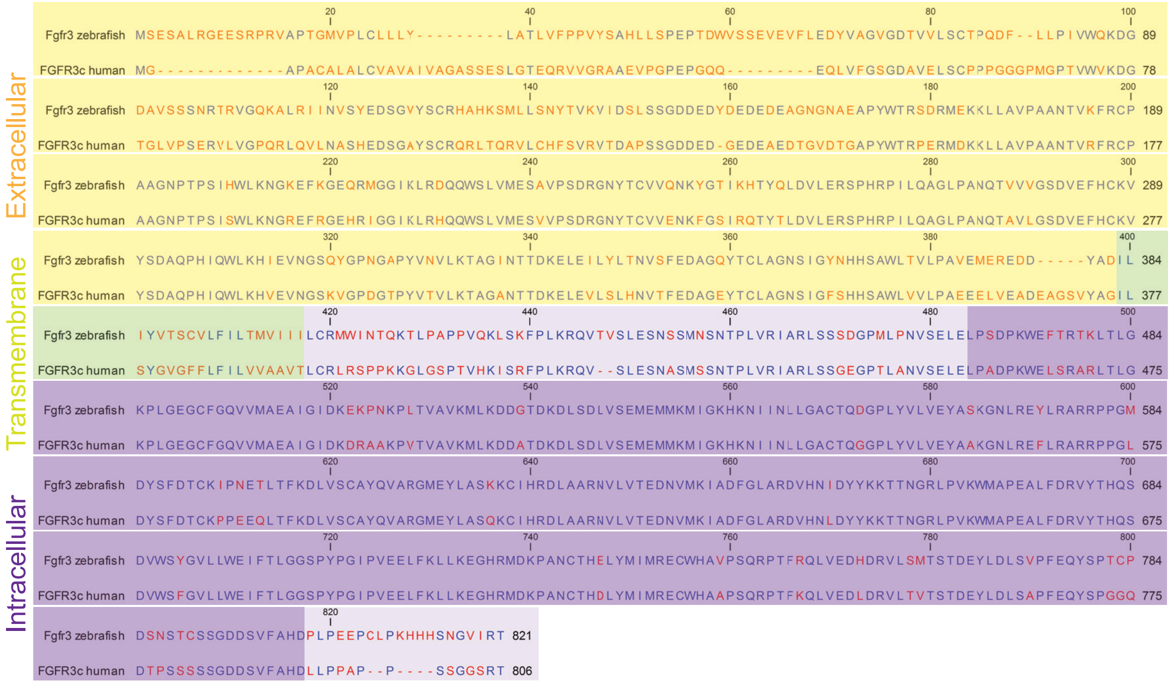

**Figure 1 supplement 1: Fgfr3's amino acid sequence is highly conserved between zebrafish and humans.**

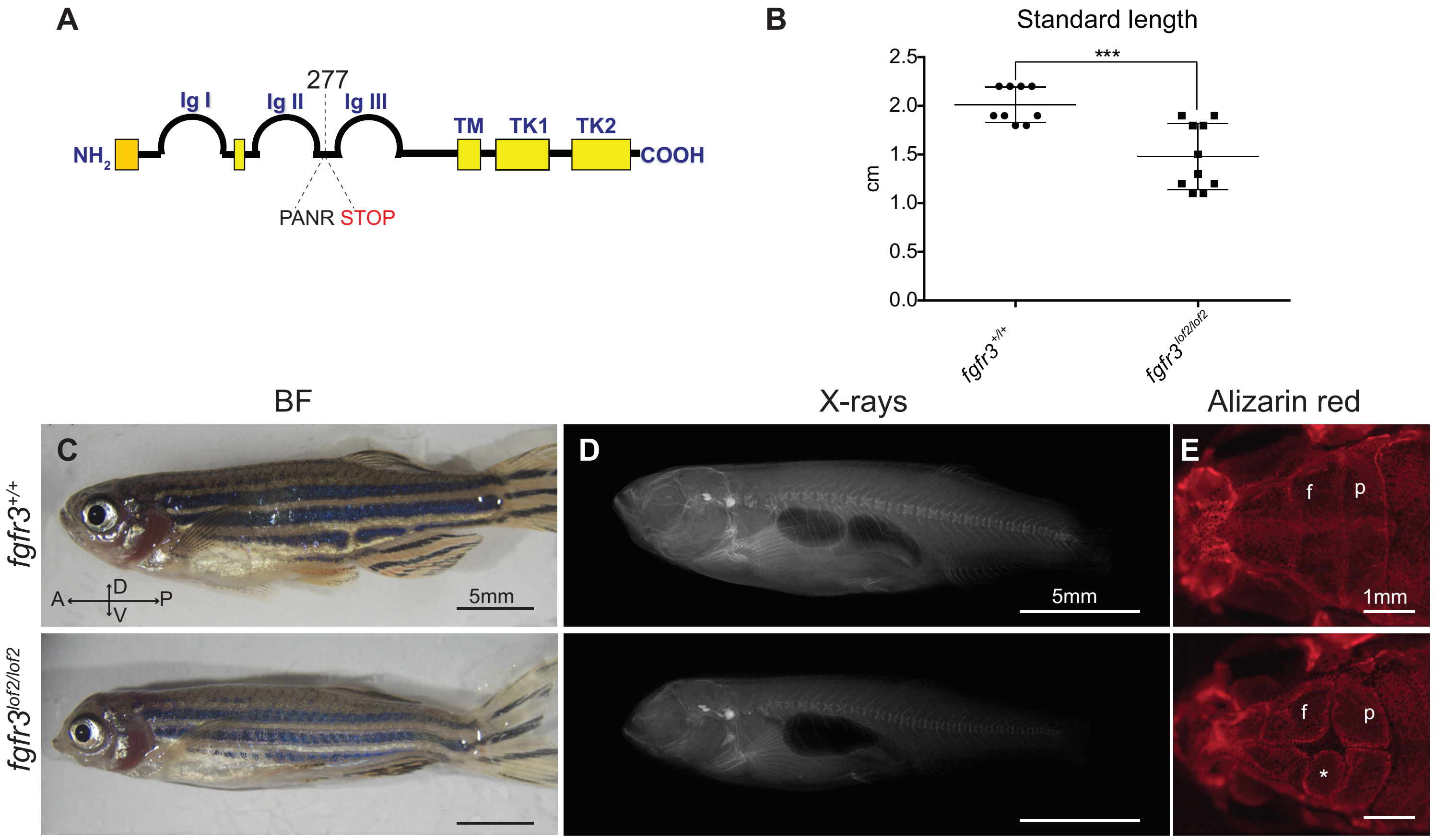


**Figure 2 supplement 1: Three-month-old *fgfr3$^{lof2/lof2}$ fish* present craniofacial and axial skeletal anomalies**

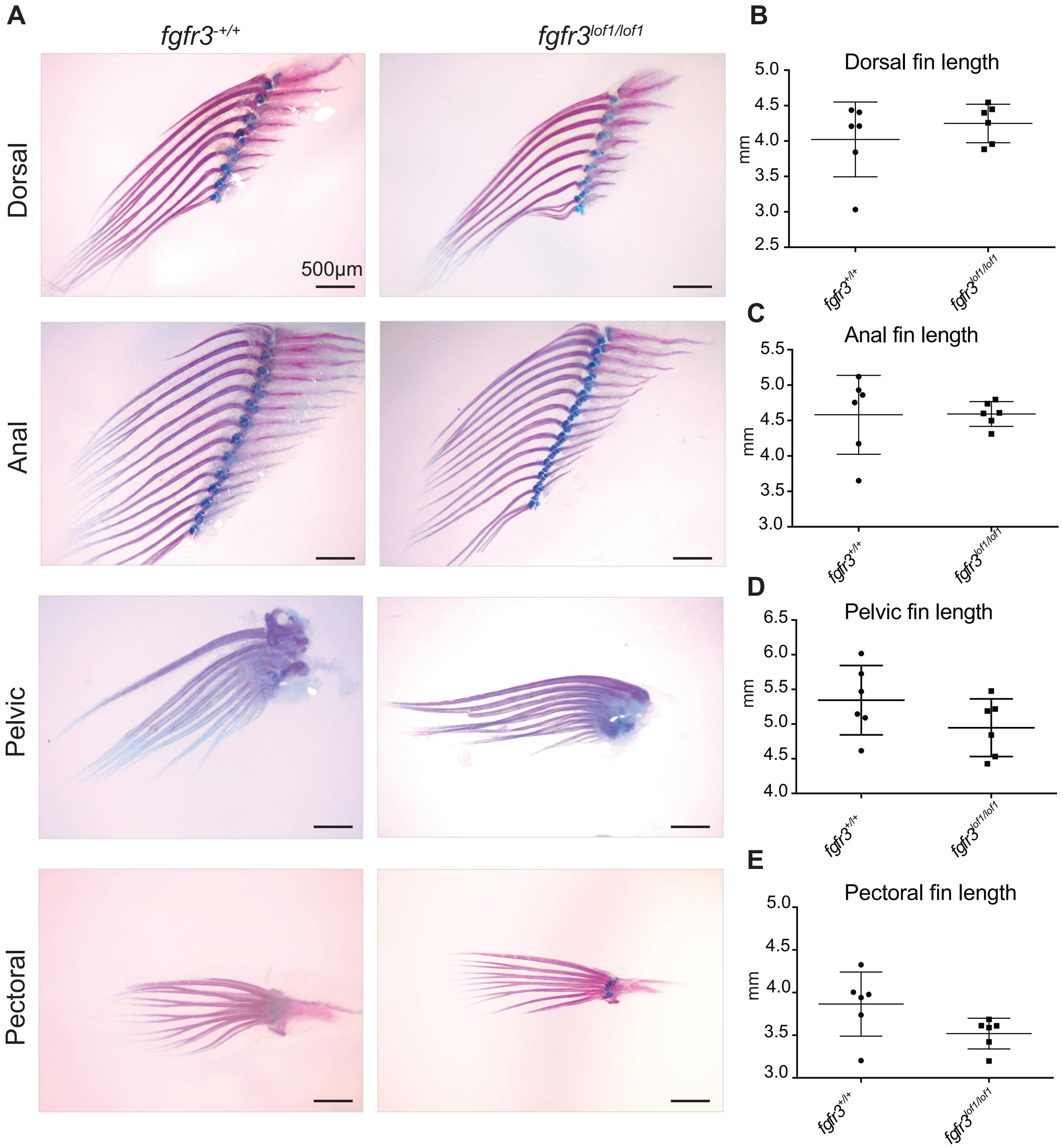


**Figure 2 supplement 2: *fgfr3* loss of function did not affect the appendicular skeleton**

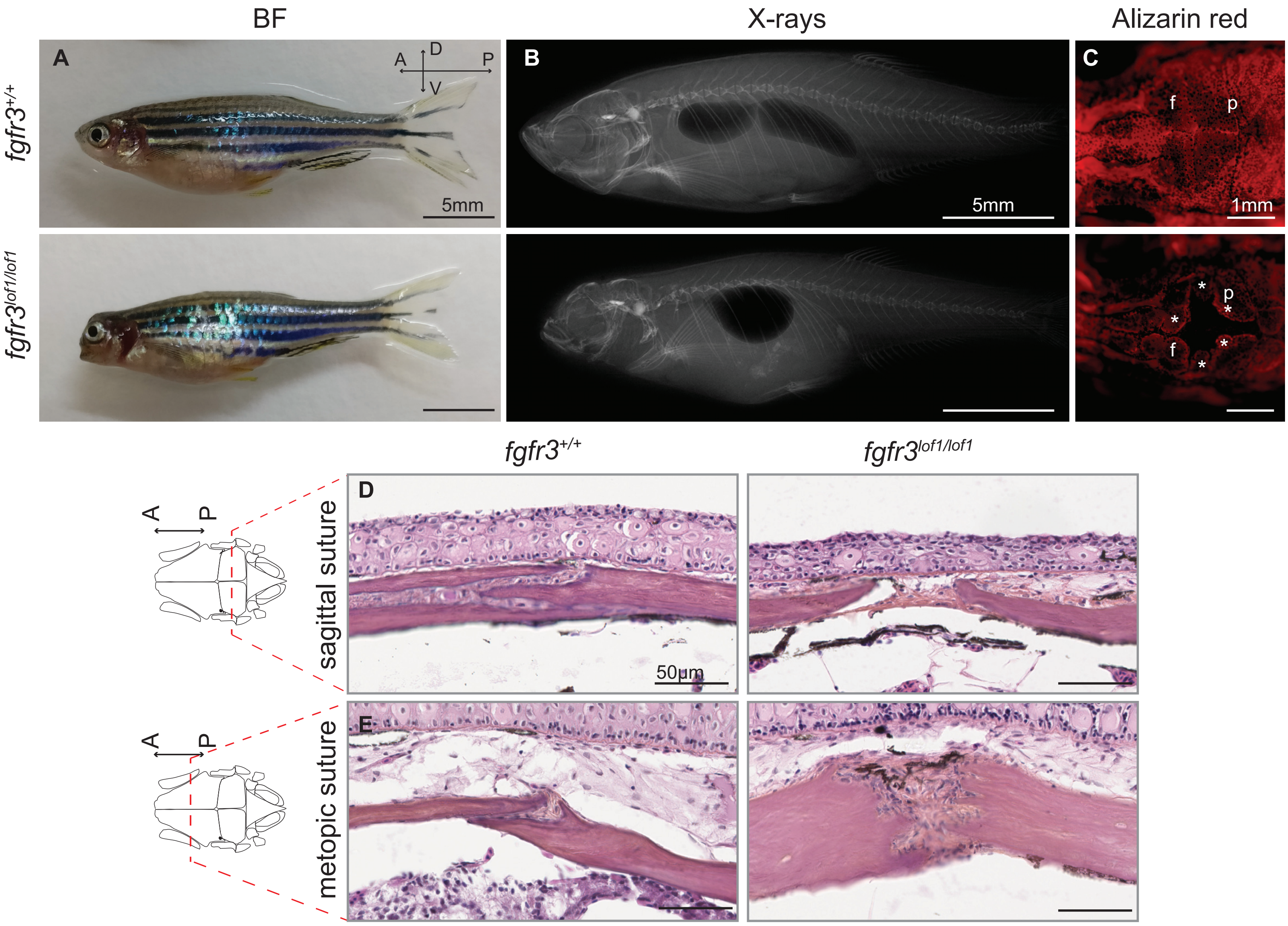


**Figure 3 supplement 1: Analyses of the skeletal phenotype in 6–month-old *fgfr3*$^{lof1/lof1}$ fish**

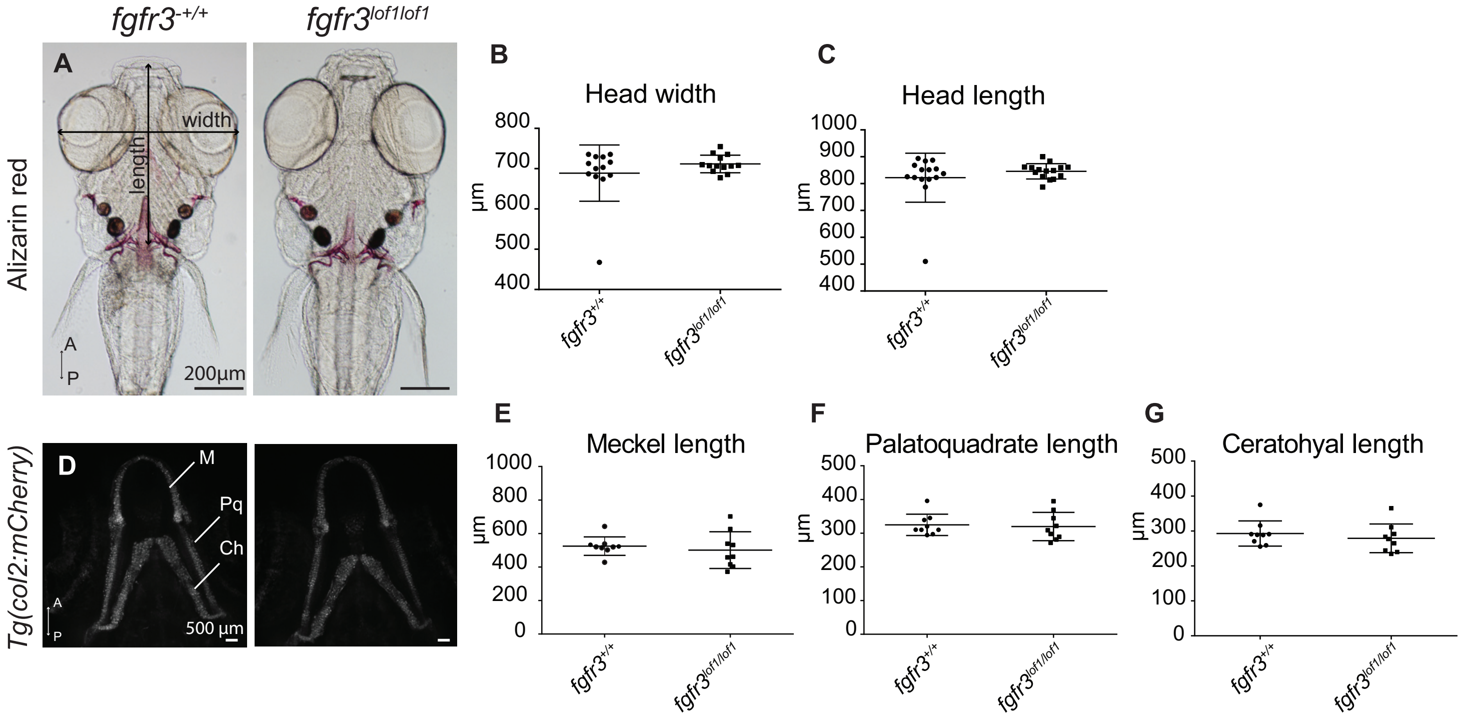


**Figure 3 supplement 2: Analyses of the larval *fgfr3*$^{lof1/lof1}$ viscerocranium**

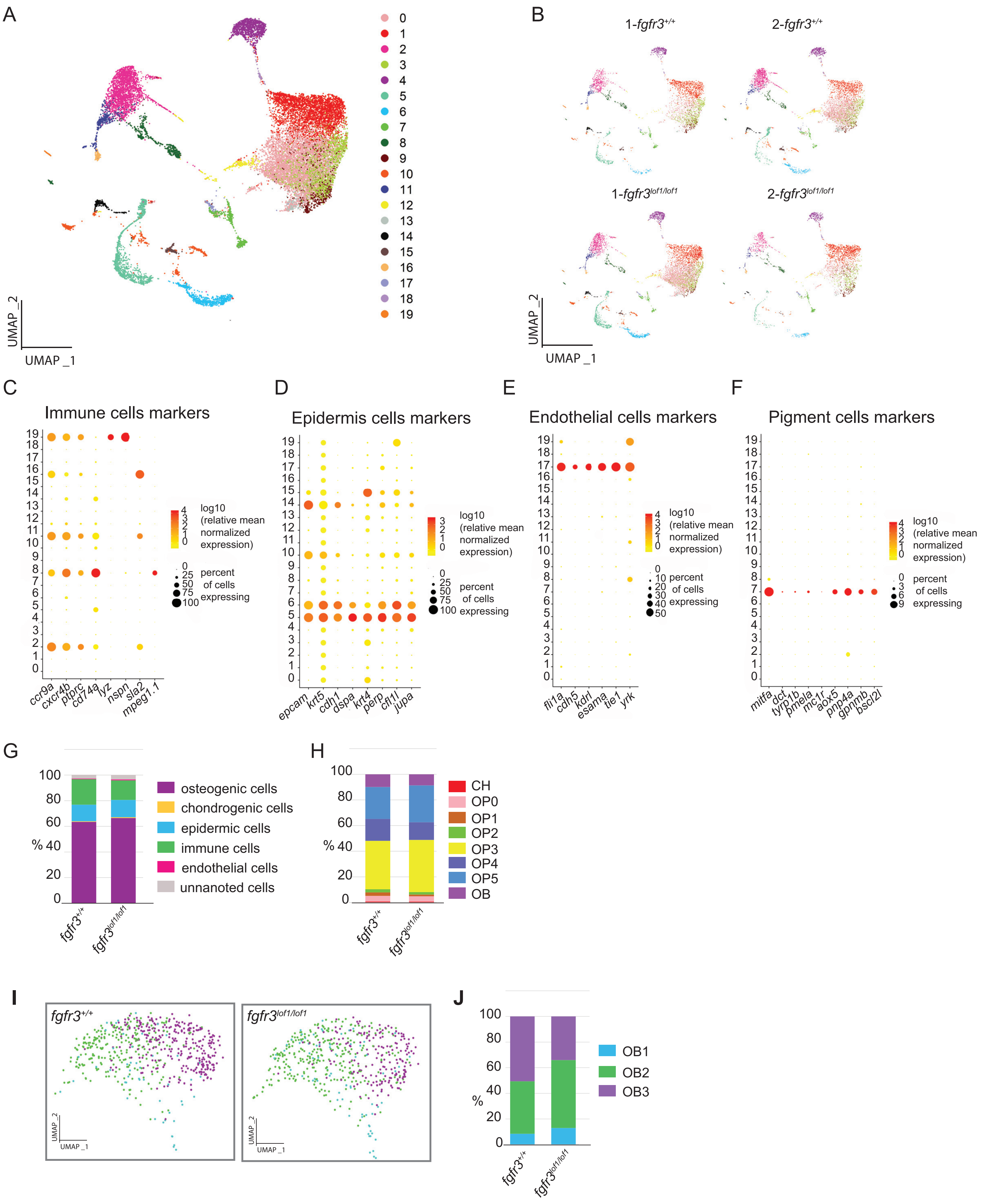


**Figure 6 supplement 1: Subpopulations of osteogenic cells involving in cranial vault formation, as revealed by single-cell RNA sequencing**
